# Identifying novel links between cardiovascular disease and insomnia by *Drosophila* modeling of genes from a pleiotropic GWAS locus

**DOI:** 10.1101/2023.07.24.550429

**Authors:** Farah Abou Daya, Torrey Mandigo, Lily Ober, Dev Patel, Matthew Maher, Suraj Math, Cynthia Tchio, James A. Walker, Richa Saxena, Girish C. Melkani

## Abstract

Insomnia symptoms have been associated with cardiovascular disease (CVD), doubling the risk of incident CVD, but specific shared pathways remain poorly understood. Recently, genome-wide association studies (GWAS) identified genetic loci significantly associated with insomnia symptoms, including one locus (near *ATP5G1*, *UBE2Z*, *SNF8*, *IGF2BP1*, and *GIP*) that was previously linked with CVD in an independent GWAS. To evaluate the cell-autonomous role of genes within the 17q21 insomnia and CVD locus, we used *Drosophila melanogaster* models to perform tissue-specific RNAi knockdown of four conserved orthologues (*ATPSynC, Lsn, Bruce*, and *Imp*) in neurons and in the heart. To identify non-cell-autonomous mechanisms, we also assessed heart function in flies with neuronal-specific knockdown and sleep in flies with heart-specific knockdown. Neuronal and cardiac-specific RNAi knockdown of several of the genes conserved in *Drosophila* led to compromised sleep quality and impaired cardiac performance. Neuronal-specific knockdown of *ATPSynC, Imp, and Lsn* led to disruptions in sleep quantity and quality. Knockdown of *ATPSynC and Lsn in the heart* led to significantly reduced cardiac performance without and with cardiac dilation, respectively. Furthermore, *Lsn* and *ATPSynC-*suppressed hearts showed disruption in the actin-containing myofibrillar organization and led to a significantly shortened lifespan. Non-cell-autonomous effects were seen both from neurons to heart (*Imp*), and heart to neurons (*ATPSynC* and *Lsn)*. Specifically, *Imp* neuronal knockdown led to a significantly compromised cardiac function, whereas knockdown of *ATPSynC* and *Lsn* in the heart led to compromised sleep characterized by increased sleep fragmentation, both accompanied by an increase in inflammation through Upd3, an inflammatory cytokine, in the heart or head, respectively. We also demonstrate disrupted cardiac function or sleep upon cardiac-specific or neuronal-specific overexpression of Upd3, respectively, showing a direct link between cardiac dysfunction and sleep disruption through inflammation. Our study reveals tissue-specific and cross-tissue consequences of Drosophila knockdown of multiple genes at this locus, providing novel insights into potential genetic mechanisms linking CVD and insomnia. Our study also highlights the key role of these four conserved genes in both sleep and cardiac function.

## Introduction

Cardiovascular disease (CVD), one of the leading causes of death worldwide, encompasses several conditions that affect heart structure and function.^1–3^ The incidence of CVD continues to rise, with approximately 18.2 million deaths worldwide in 2019, which contributes to rising healthcare costs and creates a significant socioeconomic burden.^3,4^ Many factors that increase the risk of CVD include genetic factors, smoking, and lack of physical activity. One important risk factor for CVD that has recently emerged is sleep dysfunction including insomnia.^4–7^ Insomnia is the most common sleep disorder, affecting 10 to 30% of the population, and is defined as persistent difficulty in falling and/or staying asleep or non-restorative sleep resulting in daytime sleepiness, fatigue, or dysfunction.^8–10^ Studies suggest that insomnia has a genetic component with heritability estimates ranging between 22 and 25% in adults^11,12^, and multiple GWASs have identified genetic loci with links to insomnia.^12–15^ Although genetic factors have been identified as contributors to CVD and insomnia, the genetic mechanisms underlying these two diseases remain poorly understood.

Observational studies have demonstrated that insomnia increases the risk of several disorders, especially CVD.^16–19^ Moreover, mendelian randomization analyses show that insomnia symptoms double the risk for incident CVD.^12,13^ Similarly, cardiac dysfunction has been associated with sleep disruptions.^20,21^ Furthermore, a recent study found that sleep modifies atherosclerosis through hematopoiesis in mice.^22^ Together these findings establish a clear link between cardiovascular traits and sleep disruptions. However, the specific underlying causal genetic pathways and mechanisms connecting CVD and insomnia are unknown. Recent genome-wide association studies (GWASs) identified multiple significant loci for self-reported insomnia symptoms in UK Biobank and 23andMe participants.^12,13^ From these loci, we identified a single locus, represented by lead SNP rs4643373, that has also been previously associated with coronary artery disease (CAD), and other cardiac disorders, including myocardial infarction.^23–27^ This locus provides a valuable opportunity to identify genes important for CVD and/or insomnia, and dissect potential genetic mechanisms underlying the link between cardiovascular function and sleep. Near this locus, we identified five candidate genes, *ATP5G1*, *UBE2Z*, *SNF8*, *IGF2BP1*, and *GIP*. The known functions of these genes are very diverse, including energy metabolism (ATP5G1), protein ubiquitination (UBE2Z), multivesicular body biogenesis (SNF8), post-transcriptional regulation (IGF2BP1), and lipid metabolism (GIP).^28–32^ However, it remains unclear which of these genes, if any, contribute to CVD or insomnia.

To elucidate the impact of these candidate genes on the regulation of cardiac function and sleep, we identified conserved *Drosophila melanogaster* orthologs for the insomnia and CVD-related candidate genes near rs4643373*. Drosophila* genes, namely *ATPSynC (CG1746), Bruce (CG6303), Lsn (CG6637) and Imp (CG1691),* were identified as orthologs of *ATP5G1*, *UBE2Z*, *SNF8*, and *IGF2BP1,* respectively, however, GIP doesn’t have an ortholog*. Drosophila* has become a well-established model system for studying both CVD and sleep disturbances.^33–38^ The fly heart displays many developmental and functional similarities to the mammalian heart.^33,39^ Moreover, several genes causing heart disease in humans are present in *Drosophila* and play similar pathophysiological roles, and the manipulation of these genes in *Drosophila* leads to disease phenotypes similar to humans.^33,39,40^ Sleep in flies has also been demonstrated to share many characteristics with human sleep, such as consolidation during the night and similar responses to sleep altering drugs.^34,41–43^ Therefore, studies investigating the role of human-relevant *Drosophila* orthologs in the regulation of cardiovascular function and sleep provide an efficient means to identify new causal genes related to CVD and/or insomnia, and understand mechanisms relating both diseases to identify potential future therapeutic targets.

Here, we have evaluated the cardiac and sleep roles of *Drosophila* genes *ATPSynC, Bruce, Lsn and Imp,* in both, cell-autonomous and non-cell-autonomous manners. To assess the role of these genes in cardiac and sleep physiology, we performed tissue specific knockdown (KD) in the heart and nervous system, respectively. Cardiac- and neuronal-specific KD of these genes led to cardiac and sleep dysfunction, suggesting tissue-specific functions related to each disease. After characterizing the cell-autonomous role of *ATPSynC*, *Bruce*, *Lsn*, and *Imp* in cardiac function and sleep, we also identified non-cell-autonomous effects of these genes on cardiac and sleep phenotypes, upon KD of these genes neuronally or in cardiac tissue, respectively. Neuronal KD of *Imp* compromised cardiac function, while cardiac KD of *ATPSynC, Lsn and Bruce* led to sleep dysfunction in a non-cell-autonomous manner. We further identified inflammation as an underlying mechanism involved in the non-cell-autonomous effects of cardiac dysfunction on sleep disruption and vice versa. In conclusion, we were able to uncover novel genetic mechanisms with cell-autonomous effects on the regulation of cardiac function and sleep as well as non-cell autonomous genetic mechanisms linking cardiac function in the regulation of sleep and sleep on cardiac function. Taken together, our data are among the first functional genetic proofs, to our knowledge, that link CVD with sleep disorders and provide mechanistic insight into potential therapeutic targets to prevent or attenuate both diseases.

## Results

### Shared GWAS locus at 17q21 is associated with CAD/CVD and insomnia

A GWAS signal at 17q21.32 for CAD including myocardial infarction, percutaneous transluminal coronary angioplasty, coronary artery bypass grafting, angina or chronic ischemic heart disease from the CARDIoGRAMplusC4D Consortium, represented by lead SNP rs4643373 (Fig. 1A, n=181,522 cases and 984,079 controls; OR=1.04 (95% CI=1.028-1.050))^44^ was found to co-localize with an association signal for insomnia symptoms (n=593,724 cases and 1,771,286 controls; OR=1.04 (95% CI=1.025-1.046); Fig. 1B; pp = 0.95 that both traits share the same causal SNP)^45^. The genomic region around the co-localized signal encompasses multiple genes including *ATP5MC1, UBE2Z, SNF8, GIP and IGF2BP1*. Multi-tissue eQTL analyses shows an effect of lead SNP rs4643373 on the expression of these genes in multiple tissues including the left ventricle of the heart and the hypothalamus (Table S1). Moreover, to check for other CVD-related signals in the region of rs4643373, we catalogued cardiovascular and cardiometabolic trait associations at genome-wide significance (p<5×10^-8^) of this region from literature and the Cardiovascular Disease Knowledge Portal (see methods). We found a total of 9 significant associations with cardiovascular and cardiometabolic traits, including myocardial infarction, diastolic blood pressure, and triglycerides (Fig. 1C) reinforcing the importance of this genomic region in CVD. The causal genes and variants at this locus are unknown. Furthermore, it is unclear if the association signals reflect independent contribution of effector genes to sleep and cardiovascular disease, or if effector genes influence sleep through cardiovascular dysfunction or vice versa. Thus, we developed a *Drosophila* screening approach to identify the role of fly orthologs of these genes in sleep and cardiovascular function in both cell-autonomous and non-cell-autonomous manners (Fig. 1D).

**Fig. 1.**
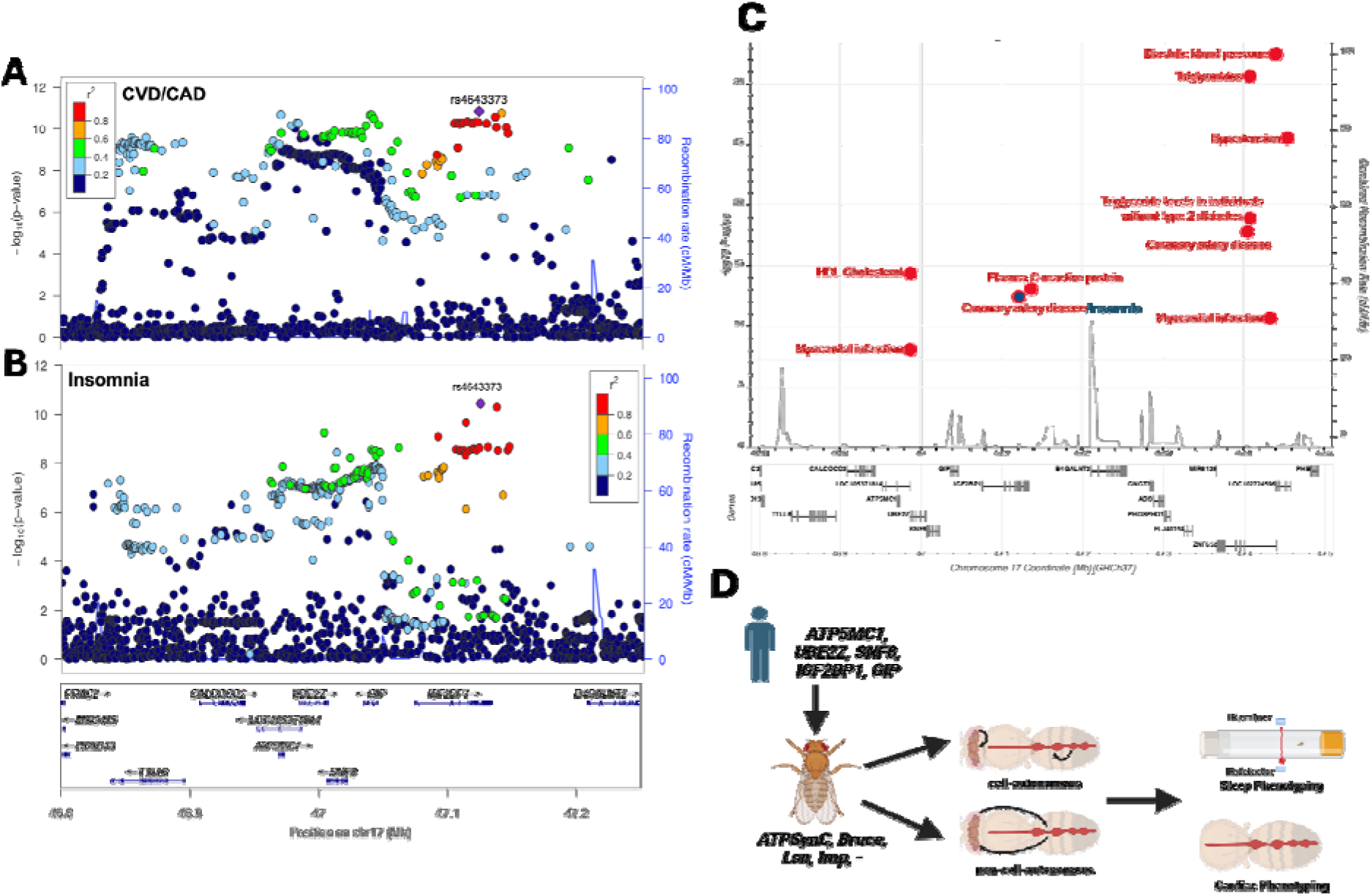
CVD- and insomnia-related locus, other phenotypic associations and nearby genes. Manhattan plots (LocusZoom) showing CVD (A) and insomnia (B) SNP association peaks with 5 nearby candidate genes, *ATP5MC1 (ATP5G1), UBE2Z, SNF8, GIP, IGF2BP1*. GWAS associations meeting genome-wide significance (p<^5×10–8^, blue line) for cardiovascular/cardiometabolic traits in the region near rs4643373 obtained from the Cardiovascular Disease Knowledge Portal (see methods) and literature. Each point corresponds to a single trait in a single study. Blue dot represents rs4643373 (C). Graphical Scheme showing workflow (D).

### Neuronal-specific suppression of CVD- and insomnia-related genes leads to altered sleep phenotypes along with enhanced sleep fragmentation

Fly orthologs of four of the five human genes near rs4643373 with their % similarity are shown in Table 1. The fifth gene, GIP, lacks a fly ortholog. In order to assess whether any of the orthologs of the genes within the CVD- and insomnia-associated locus were essential in *Drosophila*, we performed ubiquitous KD of these genes driven by *Act5C-Gal4* driver. Ubiquitous KD of *ATPSynC* and *Lsn* led to lethality, while that of *Bruce* and *Imp* did not affect viability. We then performed a pan-muscle *KD* using the *24b-Gal4* driver to determine if the function of these genes was essential in all muscle tissues. As with ubiquitous KD, we found that pan-muscle KD of *ATPSynC* and *Lsn* resulted in lethality, whereas *Bruce* and *Imp* were viable. We also tested viability with pan-neuronal KD using the *Elav-Gal4* driver. Only *ATPSynC* Line 1 (see methods) led to lethality when suppressed pan-neuronally, suggesting an essential role for *ATPSynC* in neuronal function.

**Table 1.** Human and fly symbols of CVD- and insomnia-related genes with %similarity at a single locus.

| Human | Drosophila | % |
| --- | --- | --- |
| Symbol | Symbol | Similarity |
| ATP5G1 | ATPSynC | 83 |
| UBE2Z | Bruce | 54 |
| SNF8 | Lsn | 71 |
| IGF2BP1 | Imp | 58 |
| GIP | - | - |

To test the impact of these 4 genes near the CVD- and insomnia-associated locus on sleep, we used the neuronal-specific *Elav-Gal4* driver to KD gene expression. Level of KD of each gene in the head is shown in Fig. S1A-D. We quantified total, day and night measures of sleep, locomotor activity, sleep fragmentation, sleep bout length and sleep bout number of three-to-four-day-old male flies, averaged over a 5-day period, with pan-neuronal *Elav* KD of each of the four *Drosophila* orthologs.

Since KD of *ATPSynC* using RNAi Line 1 caused lethality, we used another available line with efficient KD when driven by *Elav-Gal4* (Fig. S1A, see methods). Compared to driver and UAS control flies, RNAi-mediated inhibition of *ATPSynC* significantly increased overall sleep duration which primarily resulted from an increase in nighttime sleep (Fig. 2A-D, Table S2). This increased sleep corresponded to a decrease in overall locomotor activity (Fig. 2E, Table S2). KD of *Imp* also showed increased sleep which was primarily due to increased daytime sleep (Fig. 2A-D, Table S2) with decreased activity (Fig. 2E, Table S2). Even though the suppression of *Lsn* showed a significant increase in nighttime sleep (Fig. 2A-D), it was not significantly different than its UAS control (Table S2). However, neuronal KD of *Bruce* decreased daytime sleep but had no effect on overall sleep or activity (Fig. 2A-E, Table S2). We observed similar sleep and activity trends in *ATPSynC*, *Lsn* and *Bruce* females, but no change in *Imp* females compared to controls (Fig. S3A). Since we observed a decrease in activity accompanying an increase in sleep, we evaluated fly locomotion speed using MARGO.^46,47^ In males, we observed similar trends in locomotion speed as those measured using the DAM system in *ATPSynC* and *Imp* males (Fig. S4A). However, *ATPSynC* females, which also showed an increase in sleep, did not significantly affect locomotion speed measured using MARGO (Fig. S4B). This suggests that the increase in sleep in *ATPSynC* flies is not only due to a decrease in locomotor activity, however, the increase in sleep in *Imp* flies could be due to an overall decrease in fly activity.

**Fig. 2.**
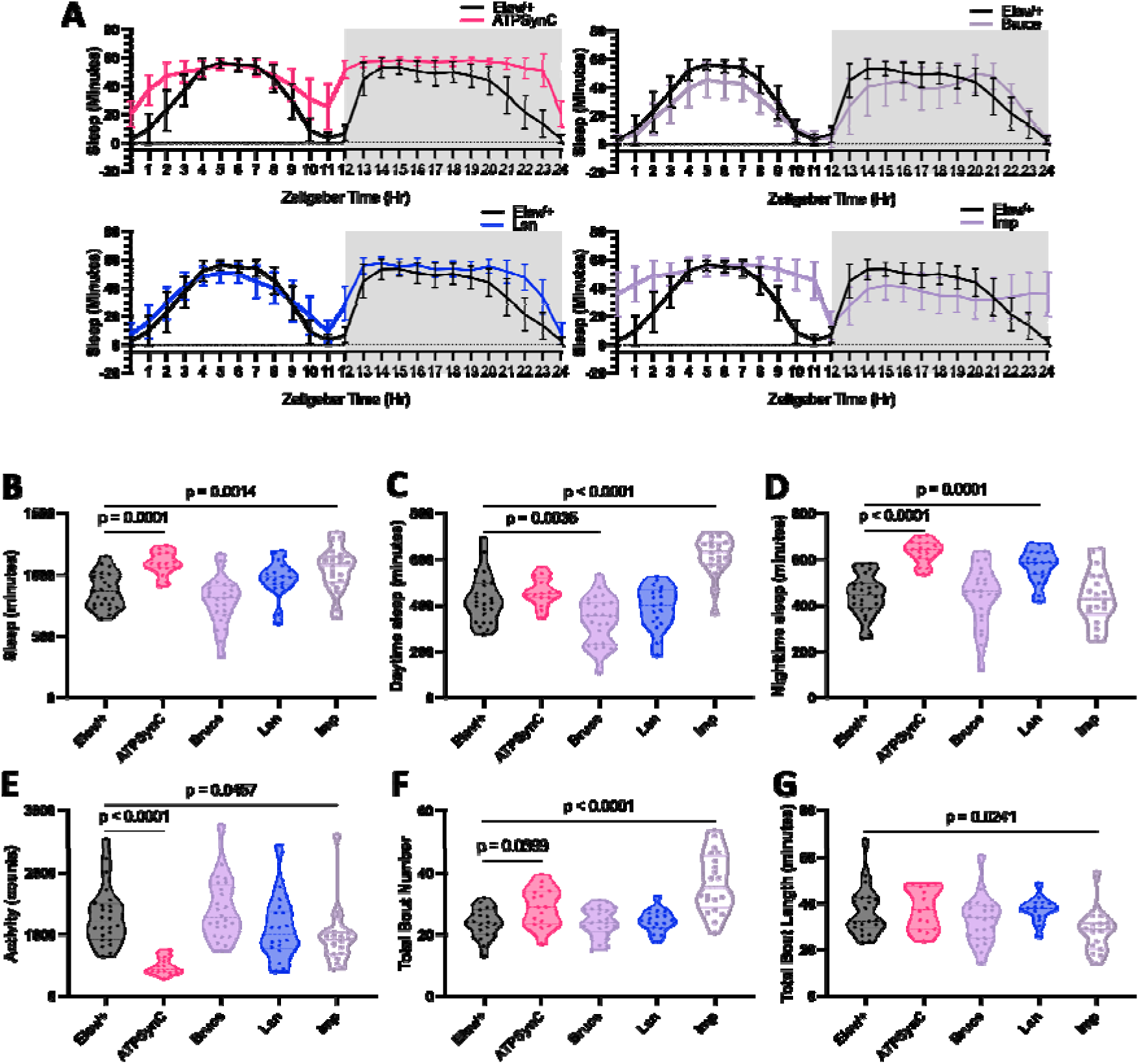
Neuronal-specific suppression of CVD- and insomnia-related genes leads to compromised sleep phenotypes. Sleep profiles showing sleep minutes per hour for 24 hours (A). Violon plots for quantitative sleep parameters, total sleep amount (B), daytime sleep amount (C), nighttime sleep amount (D), total locomotor activity (E), total bout number (F), and total bout length (G) from 1-week-old male *Drosophila* with neuronal RNAi knockdown of CVD- and insomnia-related genes with the pan-neuronal *Elav-Gal4* driver. N=16-24 for each group. For *ATPSynC*, Line 1 was lethal; Line 2 was used (refer to methods section). Data was collected from at least 2 independent experiments from one RNAi line (second line data shown in Fig. S2). Each data point represents a fly. Statistics were calculated by one-way ANOVA for comparisons to controls.

To further characterize these sleep alterations observed upon KD of each gene, we measured the number and length of sleep bouts to assess sleep fragmentation. Compared to driver and UAS controls, KD of only *Imp* resulted in increased sleep bout number along with a decreased bout length which are indicative of sleep fragmentation (Fig. 2F, G, Table S2). We also observed similar sleep and activity trends in 3-week-old male flies (Fig. S5A). Moreover, to assess the effects of adult-specific KD of those genes, we used the GeneSwitch system, and observed a consolidated increased sleep phenotype in *ATPSynC* flies similar to that observed when the gene was KD throughout development indicating the importance of this gene for sleep regulation, while sleep dysfunction observed upon *Imp* KD was lost with adult-specific KD of this gene (Fig. S6A). To determine the effect of KD of these genes on circadian rhythms, we measured rhythmicity of flies under constant darkness. We found that neuronal-specific KD of *Imp* led to circadian defects as all flies were arrhythmic (Table S3). KD of *ATPSynC* decreased the number of rhythmic flies (41.18%) but it did not significantly affect the period length or rhythm strength (FFT), while KD of *Lsn* or *Bruce* did not affect rhythmicity (Table S3). Therefore, the neuronal suppression of genes within the CVD- and insomnia-related locus led to a significantly altered sleep phenotype characterized by an increase in overall sleep in two genes with only *Imp* affecting sleep fragmentation and rhythmicity.

### Cardiac-specific suppression of CVD- and insomnia-related genes leads to cardiac dysfunction, myofibrillar disorganization, cardiac fibrosis, and shortened lifespan

To evaluate the effect of suppressing these genes on cardiac performance, KD of *ATPSynC*, *Bruce*, *Lsn, or Imp,* was carried out using the cardiac-specific *Hand-Gal4* driver. Levels of KD of each gene in the heart is shown in Fig S1E-H. One-week-old male and female flies were dissected and imaged for assessment of cardiac physiological parameters. Interestingly, suppressing *Lsn* led to a non-beating, heart failure-like phenotype where only 62.75% of hearts beat at 1 week of age which decreased to 15.91% by 3 weeks of age in males (Fig. 3A). *ATPSynC* flies also exhibited a decrease in the number of beating hearts with age, from 91.67% at 1-week to 60% beating hearts at 3-weeks of age (Fig. 3A). This suggests an important role for *Lsn* and *ATPSynC* in cardiac function. Upon analyzing beating hearts, cardiac-specific KD of *ATPSynC* showed a bradycardic phenotype with cardiac dysfunction characterized by significantly increased heart period (HP), arrhythmia index (AI), diastolic interval (DI) and systolic diameter (SD) and reduced diastolic diameter (DD) and fractional shortening (FS), a measure of cardiac performance, in both male and female flies (Fig. 3B-H, Fig. S3B, Table S2). Suppression of *Lsn* led to cardiac disfunction characterized by significantly increased DD and SD and reduced FS in both sexes, in addition to a tachycardic phenotype in males characterized by decreased HP (Fig. 3B-H, Fig. S3B, Table S2). Suppression of *Imp* led to a tachycardia characterized by decreased HP without affecting cardiac function in males only (Fig. 3B-H, Fig. S3B, Table S2). However, suppressing *Bruce* in males and females did not significantly affect heart function in one-week-old flies (Fig. 3B-H, Fig. S3B, Table S2). To assess whether or not the effects of these genes on the heart persist with age, we assessed cardiac function in 3-week-old male flies with cardiac-specific suppression of these genes and observed similar overall trends for cardiac parameters as observed in 1-week-old *ATPSynC* and *Lsn* flies (Fig. S5B). Interestingly, in 3-week-old flies with KD of *Bruce*, we observed cardiac dysfunction characterized by increased DD and SD and decreased FS (Fig. S5B), suggesting an age-related component involved in the cardiac phenotype observed with *Bruce* KD. To determine the effect of adult-specific KD of these genes, we used the GeneSwitch system. Upon adult-specific KD of *ATPSynC* or *Lsn*, we observe significantly decreased fractional shortening similar to that observed upon KD throughout development; thus, confirming that *ATPSynC* or *Lsn* are essential genes for cardiac function (Fig. S6B).

**Fig. 3.**
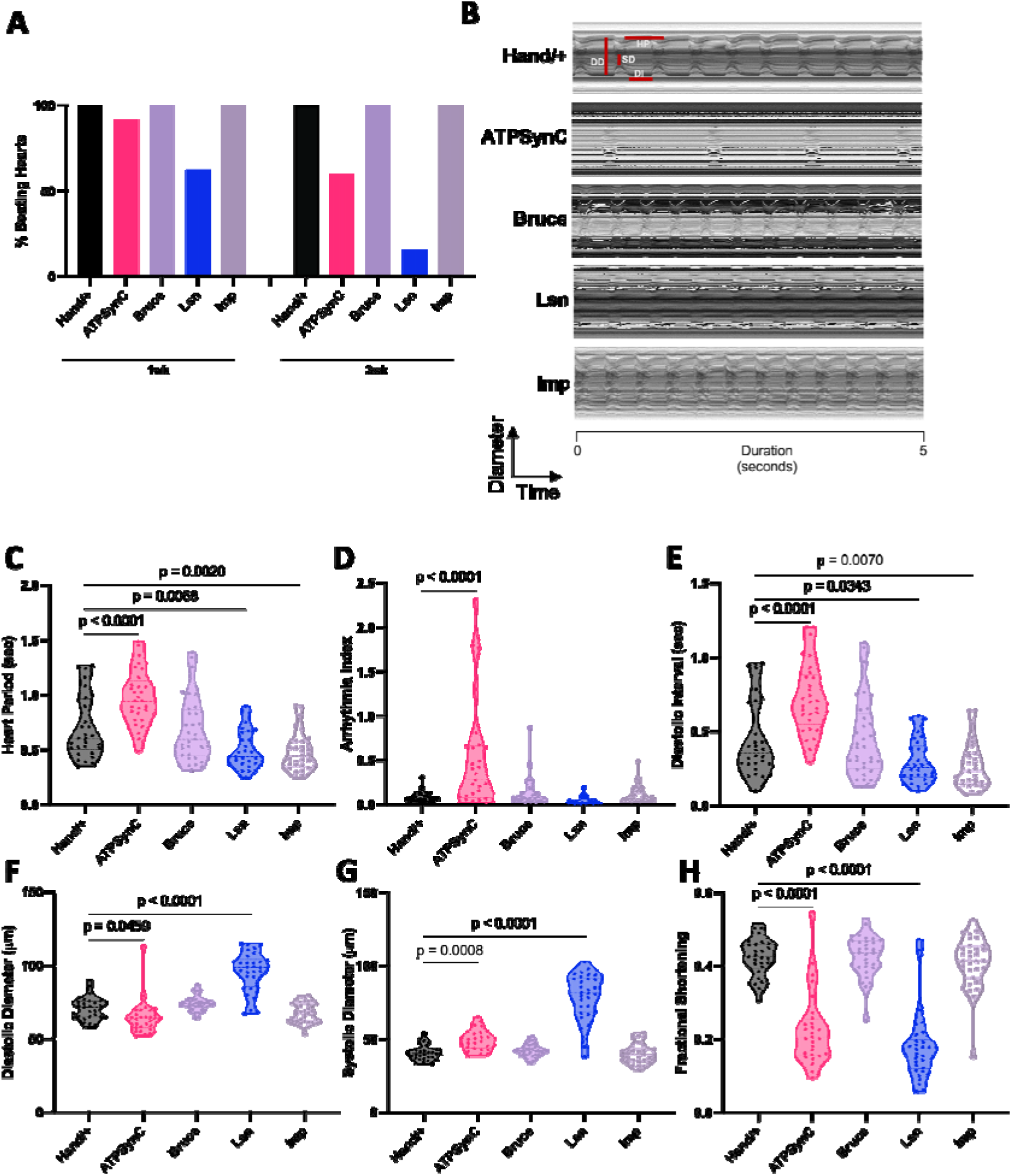
Cardiac-specific suppression of CVD- and insomnia-related genes leads to cardiac dysfunction. Representative 5-second mechanical-modes (A) from 1-week-old male flies with cardiac RNAi knockdown of CVD- and insomnia-related genes with cardiac-specific *Hand-Gal4* driver. Percentage of beating hearts at 1 versus 3 weeks of age shows significant effect of Lsn KD with age (p<0.0001) (B). Violin plots for cardiac physiological parameters, heart period (C), arrythmia index (D), diastolic interval (E), diastolic diameter (F), systolic diameter (G), fractional shortening (H), and N= 29-33 for each group for C-H, N=32-51 for each group for B. Each data point represents one fly. Data was collected from at least 2 independent experiments from one RNAi line (second line data shown in Fig. S2). Statistics were calculated by 1-way ANOVA for C-H. Fisher’s exact test was performed for A.

To characterize the morphology of hearts with KD of these genes, 1-week-old male hearts were stained with phalloidin. KD of *ATPSynC* severely disrupted actin-containing myofibrillar organization and led to almost complete loss of contractile circumferential muscles (CF) and mostly non-contractile longitudinal muscles (LF) are seen (Fig. 4A, Fig. S7A). KD of *Lsn* showed a dilated heart with more evident CF aggregations along with myofibrillar disarray, while that of *Bruce* and *Imp* showed a less severe phenotype, with visible CFs and LFs (Fig. 4A, Fig. S7A). Moreover, only suppression of *Lsn* led to significantly increased pericardin deposition, which is a collagen-like protein and a component of the extra cellular matrix (ECM) indicative of a fibrotic phenotype (Fig. 4B, C). We also assessed the effects of cardiac KD of these genes on lifespan. Cardiac suppression of *ATPSynC, Lsn,* and *Imp* led to a significantly shortened lifespan in both sexes (p< 0.0001), while flies with suppressed *Bruce* showed an increased lifespan in males (p=0.0057) (Fig. 4D). Our findings indicate that suppression of CVD- and insomnia-related genes *Lsn* and *ATPSynC* in the heart led to significantly compromised cardiac function with myofibril disorganization and shortened lifespan. Moreover, *Bruce* and *Imp* showed less severe phenotypes with *Bruce* leading to cardiac dysfunction with increased age.

**Fig. 4.**
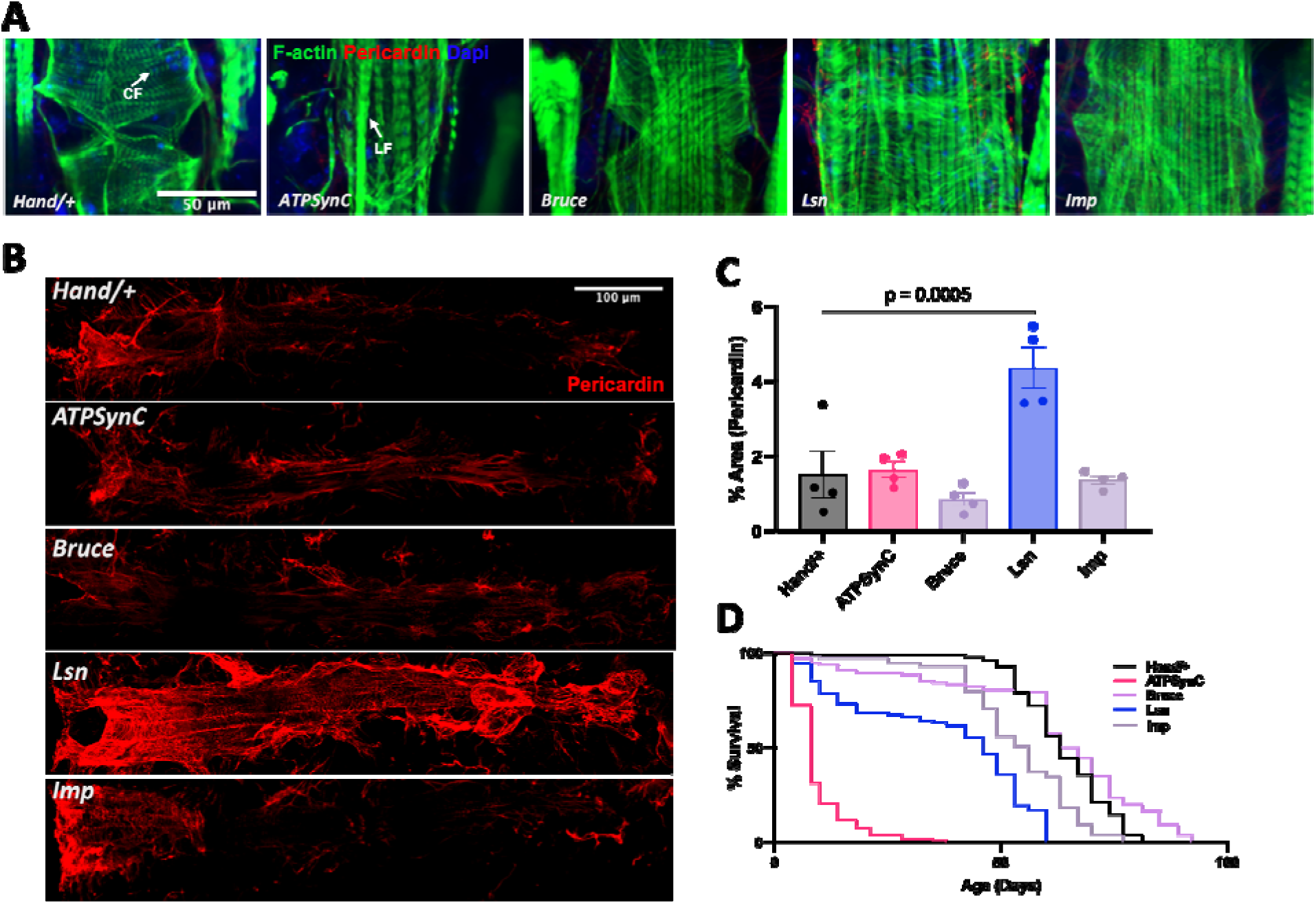
Cardiac-specific suppression of CVD- and insomnia-related genes leads to myofibrillar disorganization, cardiac fibrosis, and shortened lifespan. Representative images showing actin-containing myofibrils (A) and pericardin (B) from 1-week-old male flies with cardiac RNAi knockdown of CVD- and insomnia-related genes with *Hand-Gal4.* Quantification of pericardin signal (C). Each data point is a fly. Lifespan assay (D) for male flies with cardiac RNAi knockdown of CVD- and insomnia-related genes with cardiac-specific *Hand-Gal4* driver resulted in significant decrease in lifespan (p<0.0001) of *ATPSynC*, *Lsn* and *Imp*, and a significant increase in lifespan of *Bruce* (p=0.0057). Graph plots % survival (n >100 for each group) vs. time post-eclosion. Statistics were calculated by 1-way ANOVA for C-D and a Kaplan-Meier test was performed for E.

### Non-cell-autonomous mechanisms linking CVD with insomnia

Mendelian randomization analyses in recent genetic studies confirm a causal role for insomnia on CVD.^12^ Moreover, cardiac dysfunction has also been associated with sleep disruptions.^20,21,48^ Therefore, to assess non-cell-autonomous roles of these genes in influencing cardiac and sleep dysfunction, we suppressed genetic expression in the neurons and measured cardiac function, or we suppressed it in the heart and assessed sleep phenotypes in 3-week-old male flies (Fig. 5A). Unlike cardiac-specific KD, neuronal suppression of *ATPSynC*, *Lsn* or *Bruce* resulted in no cardiac phenotype, while neuronal suppression of *Imp* in 3-week-old male flies significantly decreased HP, DI and significantly reduced FS (Fig. 5B-D). Neuronal-specific KD of only *Imp* led to increased sleep fragmentation (Fig. 2F, G), which may be underlying the cardiac dysfunction observed in those flies. This compromised cardiac performance indicates a non-cell-autonomous effect of *Imp* in the causal role of sleep disruption on cardiac function.

**Figure 5.**
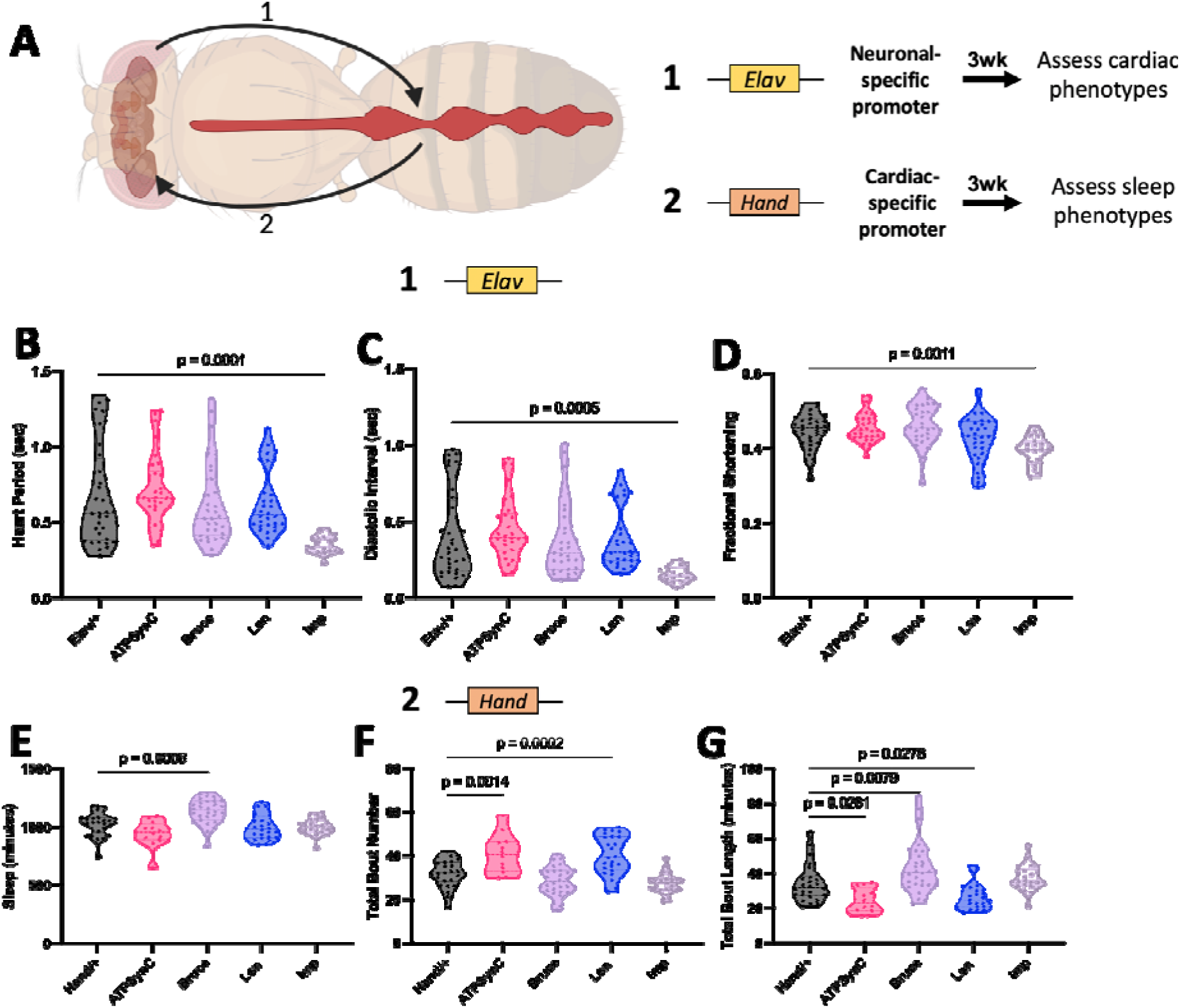
Non-cell-autonomous mechanisms linking CVD with insomnia. Graphical scheme showing experimental layout (A). Violin plots for cardiac physiological parameters, heart period (B), diastolic interval (C) and fractional shortening (D) from 3-week-old male flies with neuronal-specific knockdown of CVD- and insomnia-related genes (N= 19-32 per group). Violin plots for quantitative sleep parameters; total sleep amount (E), total bout number (F), and total bout length (G) from 3-week-old male *Drosophila* with cardiac-specific knockdown of CVD- and insomnia-related genes. N= 12-30 per group. Each data point represents one fly. Statistics were calculated by 1-way ANOVA.

Upon the cardiac-specific KD of *Bruce* in 3-week-old flies, overall sleep significantly increased (Fig. 5E). However, KD of ATPSynC or *Lsn* did not affect overall sleep (Fig.5E). We then measured the number and length of sleep bouts to assess sleep fragmentation. Only *ATPSynC* and *Lsn* showed a significant increase in total sleep bout numbers and decrease in total bout length (Fig. 5F, G) indicative of significant sleep fragmentation in these flies. In support of our data, cardiac dysfunction has been associated with sleep disruptions in observational studies.^21,48^

Since inflammation was reported as a potential mechanism underlying the connection between CVD and sleep dysfunction^18^, we measured *Upd3* levels, an IL-6-like proinflammatory cytokine in flies^49^, in heads and hearts of flies with neuronal-specific KD (Fig. 6A, B) or cardiac-specific KD (Fig. 6C, D). We first evaluated the effect of neuronal-specific KD of these genes on *Upd3* levels. *ATPSynC* KD led to increased *Upd3* levels in heads of flies while *Imp, Lsn, and Bruce* KD led to a trend towards increased Upd3 levels, but it was not significant (Fig. 6A). This may suggest that Upd3 is a mechanism involved in the sleep effects observed upon KD of these genes. Upon the KD of *Imp* neuronally, *Upd3* levels in the heart significantly increased (Fig.6B). This suggests a non-cell-autonomous effect of *Imp* in the causal role of sleep disruption on cardiac dysfunction through an increased inflammatory state. We then measured the levels of Upd3 in the heart upon cardiac-specific KD of these genes. KD of *Lsn* led to significant increase in Upd3 levels, while suppression of *ATPSynC* led to a trend towards increased Upd3 levels (p=0.0719) (Fig. 6C). This may suggest that Upd3 is the specific mechanism underlying CVD-related inflammation observed upon KD of these genes. As another measure of inflammatory-like state, we performed hemocyte counts. Cardiac-specific suppression of *Lsn* led to a significantly increased number of hemocytes in the hemolymph (Fig. S7B). Interestingly, the KD of *Lsn* in the heart led to a significant increase in *Upd3* levels in the head, while that of *ATPSynC* showed an increase that was not significant (Fig. 6D, p=0.33), both, Lsn and ATPSynC, being the only 2 genes that led to an increased sleep fragmentation phenotype non-cell-autonomously upon cardiac-specific KD (Fig. 5F,G).

**Figure 6.**
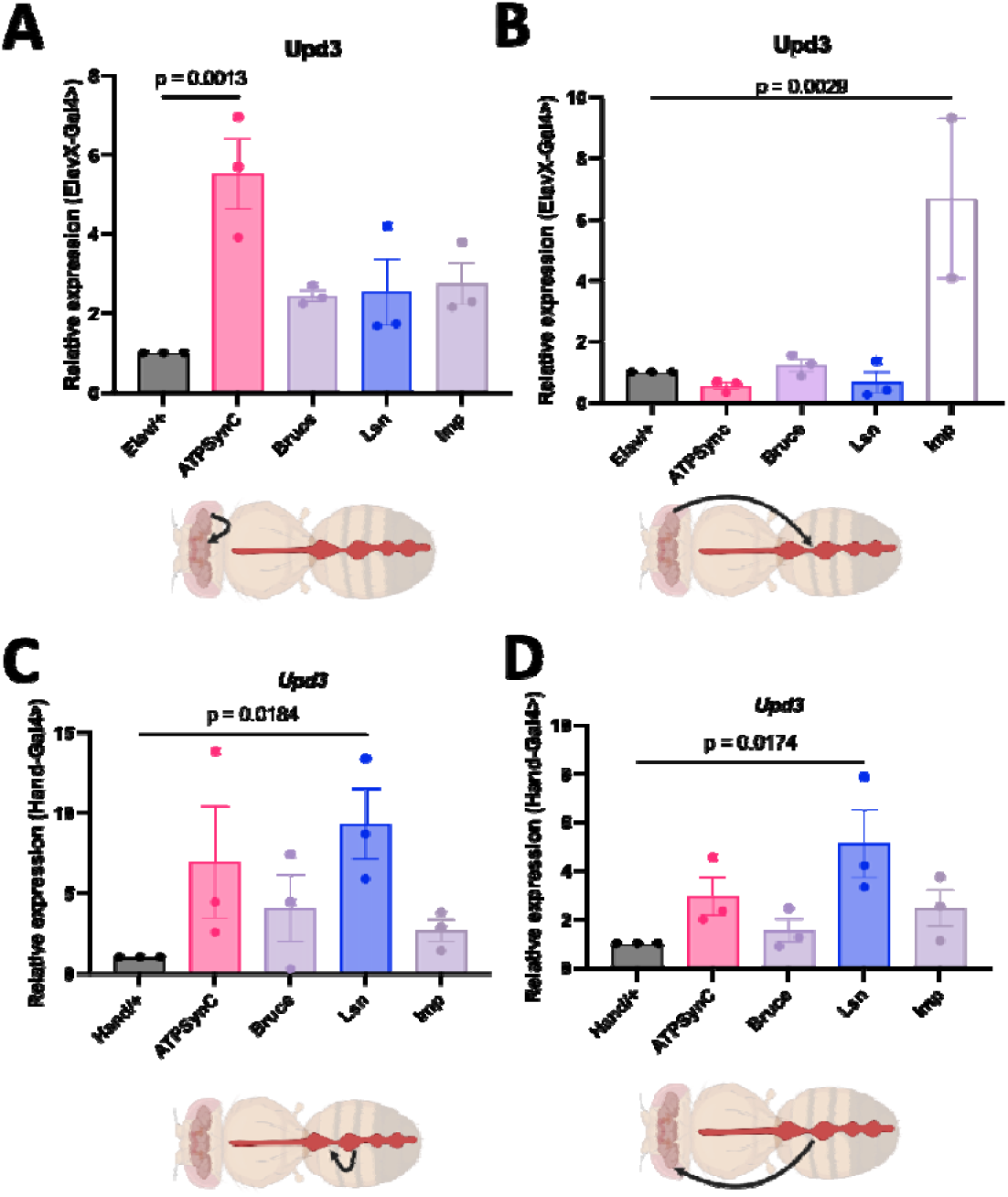
Knockdown of CVD- and insomnia-related genes increases inflammation cell-autonomously and non-cell-autonomously. Transcript levels of Upd3 in heads of flies with neuronal-specific knockdown of these genes (N= 10-12 heads per data point per group) (A). Transcript levels of Upd3 in hearts of flies with neuronal-specific knockdown of these genes (N= 10-12 hearts per data point per group) (B). Transcript levels of Upd3 in hearts of flies with cardiac-specific knockdown of these genes (N= 10-12 hearts per data point per group) (C). Transcript levels of Upd3 in heads of flies with cardiac-specific knockdown of these genes (N= 10-12 heads per data point per group) (D). Statistics were calculated by 1-way ANOVA.

To assess whether Upd3 elevation is a mechanism involved in the non-cell-autonomous effects between cardiac and sleep dysfunction, we overexpressed Upd3 in the neurons or heart using Elav- or Hand-Gal4, respectively. When we overexpressed it in neurons, it increased overall sleep amount (Fig. S8A), similar to sleep dysfunction observed in ATPSynC and Imp KD (Fig. 2B). We then assessed cardiac function upon neuronal-specific Upd3 overexpression, and we observed a tachycardiac phenotype characterized by decreased HP, DI and decreased cardiac performance reflected by decreased FS (Fig. 7A), which is similar to that observed upon the neuronal KD of Imp (Fig. 5B-D). When we overexpressed Upd3 in the heart, we observed a cardiac dysfunction phenotype similar to Lsn, characterized by increased AI, DD, and SD, and decreased FS (Fig. S8B). Interestingly, we also observed a non-cell-autonomous effect on sleep where flies with cardiac-specific overexpression of Upd3 also had significantly increased sleep fragmentation characterized by increase in bout numbers and decrease in bout lengths without effecting sleep amount (Fig. 7B) similar to the non-cell-autonomous effect of cardiac-specific KD of ATPSynC and Lsn on sleep (Fig. 5E-G). These novel findings suggest inflammation through Upd3 as an underlying mechanism linking cardiac and sleep dysfunction, bidirectionally.

**Figure 7.**
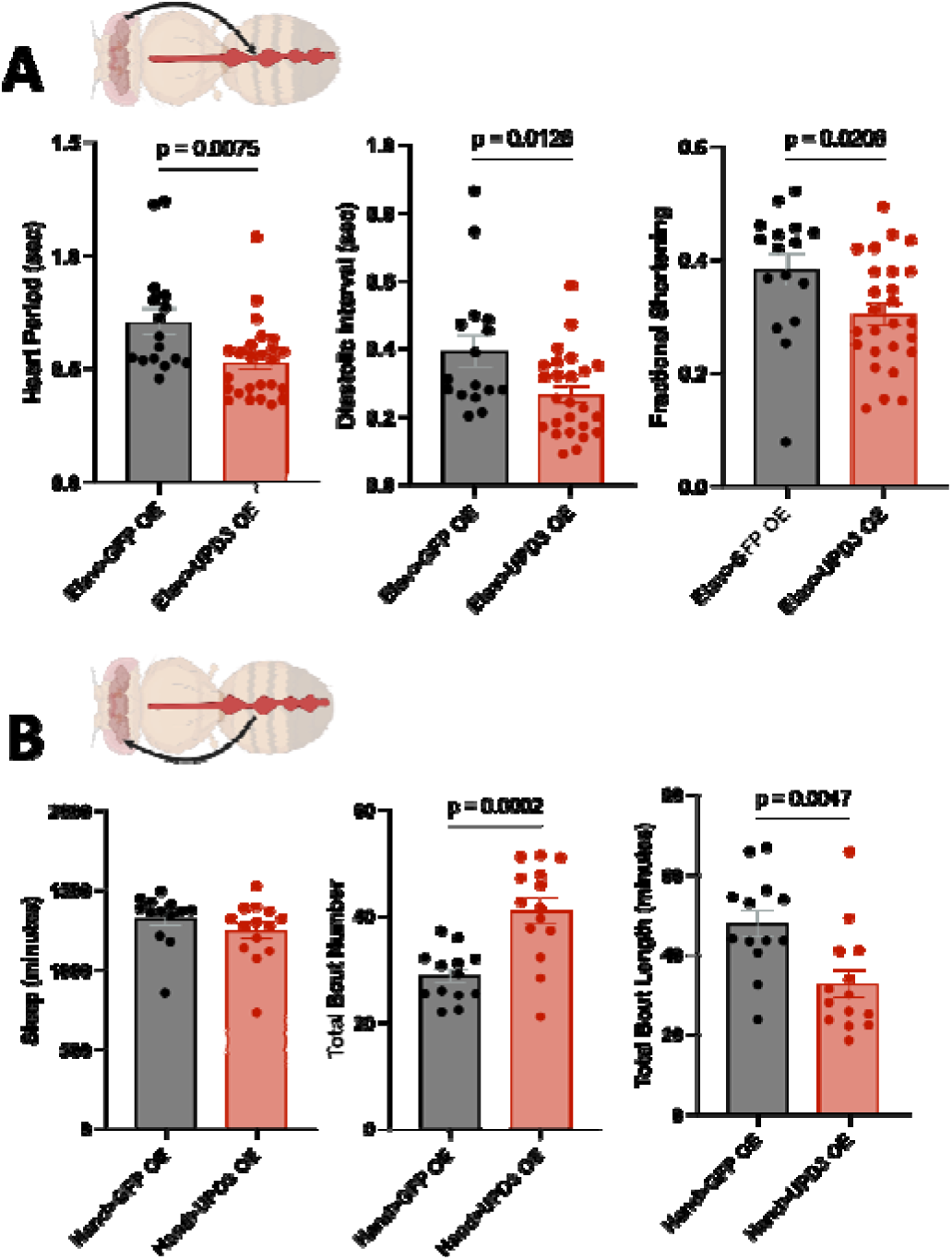
Overexpressing Upd3 in neurons or the heart leads to cardiac or sleep dysfunction, respectively. Bar graphs showing cardiac physiological parameters, heart period, diastolic interval and fractional shortening from 3-week-old male flies with neuronal-specific overexpression of Upd3 (N= 16-26 per group) (B). Bar graphs showing quantitative sleep parameters; total sleep amount, total bout number, and total bout length from 3-week-old male flies with cardiac-specific overexpression of Upd3. N= 13-14 per group. Each data point represents one fly. Statistics were calculated by unpaired t-test.

## Discussion

This study is the first to identify four genes at a single locus that link CVD and insomnia, *ATPSynC, Bruce*, *Lsn,* and *Imp* (ATP5G1, UBE2Z, SNF8, and IGF2BP1 in humans). Genetic screens have been previously applied in different model systems including zebrafish, *Drosophila* and mice to identify genes involved in different CVDs and sleep regulation.^50–56^ Despite the numerous advantages of these models for functional and behavioral screening, only few studies have utilized them to test genes identified after human GWASs. A recent study^56^ used *Drosophila* to identify causal variants reported in an insomnia GWAS^13^, including our insomnia-related locus, and screen candidate genes to pinpoint those involved in sleep regulation. However, there are no studies to date that identify genes related to both diseases or investigate functional genetic mechanisms underlying a connection between CVD and sleep dysfunction.

Here, we used an innovative approach integrating the use of human genetics in conjunction with fly genetics to identify genes related to each disease and advance the understanding of the association between CVD and insomnia. We focused on a genetic locus identified in both CVD and insomnia GWASs^13,15,23^ and identified *Drosophila* orthologs of potential nearby causal genes (Fig. 1). The locus we identified presented as a colocalization signal for both diseases. Interestingly, of 554 risk loci for insomnia identified thus far, this locus is among only three loci that colocalize with CVD (pp >0.90; others include the ApoE region, *and LINGO4/RORC*)^45^. Functional dissection of other association signals for cardiovascular/cardiometabolic traits in this same genomic region suggests an importance of this region in CVD and complex contributions of multiple genes at the locus to CVD pathogenesis.^57^

The first objective of our study was to functionally identify a novel role of these genes in CVD and/or insomnia. Thus, we performed tissue-specific, neuronal or cardiac, KD of each gene. KD of *Imp* significantly increased sleep which was fragmented and disrupted rhythmicity (Fig. 2, Table S3), thus implicating this gene with sleep regulation and maintenance of the circadian rhythm. However, KD of *Imp* also decreased the activity and speed of flies (Fig.2, Fig. S4) which could influence the increase in sleep observed in these flies. Therefore, future studies testing different levels of *Imp* KD and their effects on sleep, activity and circadian rhythms are required for a better understanding of the specific role of *Imp* on sleep physiology. KD of *Imp* did not have a severe cell-autonomous effect on the heart. Similarly, KD of *Bruce* did not affect overall sleep but started showing a decrease in cardiac function at 3 weeks of age in a cell-autonomous manner, which in turn led to a non-cell-autonomous effect on sleep duration (Fig. S5, Fig. 5).

Neuronal suppression of *ATPSynC* only increased overall sleep duration (Fig. 2). This increased sleep phenotype is supported by published findings in another study, screening insomnia-related genes identified from GWAS.^56^ This increased sleep corresponded with decreased locomotor activity, which has been recently reported in humans and flies with mutations in *ATP5G1*/*ATPsynC*^58^. However, video tracking of flies to probe for locomotion speed showed that speed was decreased only in males even though both male and female flies showed an increase in sleep amount (Fig. 2, Fig. S4) suggesting an effect of *ATPSynC* on sleep independent of locomotion defects. Neuronal KD of ATPSynC also increased inflammation (Fig. 6A), which has been associated with sleep dysfunction.^59^ Cardiac suppression of *ATPSynC* significantly compromised cardiac function characterized by severely increased arrythmia (Fig. 3) and disrupted structure and fibrosis (Fig. 4). It also increased *Upd3*-specific inflammation (Fig. 4), which is an important indicator of cardiac injury. These findings revealed a novel role of *ATPSynC* in cardiac and sleep regulation in a cell-autonomous manner. Both the neurons and heart require large amounts of ATP to perform their functions. In both organs, ATP is essential for electrophysiological activities in resting and active states^60,61^, and reduction of ATP levels impairs neural and cardiac functions^60,62,63^. ATP is produced by ATP Synthase, and impairing the function of ATP Synthase is known to be associated with cardiovascular and neurological diseases.^64,65^ *ATPSynC* is an important component of ATP Synthase. Therefore, KD of *ATPSynC* may disrupt the function of ATP Synthase thus contributing to CVD and sleep disruptions we observed.

Cardiac-specific KD of *Lsn* also resulted in a severe cardiac phenotype with significant compromised cardiac performance, characterized by significant dilation (Fig. 3), and evident myofibril disorganization and fibrosis (Fig. 4). It also showed inflammation (Fig. 6C), indicating cardiac injury, along with a unique non-beating heart phenotype that worsened with age (Fig. 3). These novel findings establish a cell-autonomous role of *Lsn* in cardiac dysfunction. *Lsn* is part of the ESCRT pathway, which is a key mechanism of multivesicular body (MVB) biogenesis.^30,66^ MVBs form exosomes which are crucial for intercellular communication and have been implicated in the pathophysiology of CVD and other diseases^30,66,67^, which supports the importance of *Lsn* for cardiac function we observed.

Our next objective was to assess associations between CVD and insomnia and assess the effects of one disease on the other through these genes (Fig. 5). First, we suppressed these genes neuronally and assessed cardiac function. Unlike cardiac KD, neuronal suppression of only *Imp* significantly reduced cardiac function which goes along the strong sleep phenotypes observed upon neuronal *Imp* KD. One possible mechanism reported to underly the effects of sleep dysfunction on cardiac function is inflammation.^18^ Notably, neuronal KD of *Imp* increased inflammation in the heart (Fig. 6B). Furthermore, the overexpression of Upd3 in neurons phenocopied those non-cell-autonomous effects on cardiac function (Fig. 7A); thus, further suggesting the influence of sleep dysfunction on cardiovascular performance through inflammation, supporting mendelian randomization reports that show an effect of insomnia on CVD.

Although previous observational and genetic studies more commonly report an effect of sleep on CVD, some human studies show an effect of heart failure on sleep interruption.^20^ Therefore, we were interested in assessing whether there is an influence in the opposing direction, from heart on the neurons. Therefore, we suppressed these genes in the heart and assessed sleep physiology (Fig. 5). We observed non-cell-autonomous effects on sleep in genes with cardiac dysfunction upon cardiac-specific KD, where flies with cardiac KD of Lsn and ATPSynC had increased sleep fragmentation. Moreover, although cardiac KD of *Bruce* did not affect heart function in 1-week-old flies, it significantly reduced cardiac function in 3-week-old flies (Fig. S5) which in turn increased sleep amount non-cell-autonomously. This shows an evident effect of cardiac dysfunction on sleep. These findings suggest a non-cell-autonomous influence of *ATPSynC*, *Lsn* and *Bruce* on sleep regulation.

While the influence of heart function on the nervous system has been reported, underlying mechanisms remain poorly understood.^68,69^ We hypothesized again that inflammation is a mechanism underlying the effects of cardiac dysfunction on sleep disruption. KD of *Lsn* and *ATPSynC* increased inflammation in the head after cardiac suppression (Fig. 6D). Importantly, we show that the overexpression of Upd3 in the heart leads to cardiac dysfunction cell-autonomously and sleep disruption non-cell-autonomously (Fig. S8B, Fig. 7B); recapitulating non-cell-autonomous effects observed by cardiac KD *ATPSynC* and *Lsn* on sleep. Overall, our results suggest inflammation as an important underlying factor in the bidirectional connection between cardiovascular function and sleep physiology.

One limitation of our study is the inability to characterize vascular and atherosclerotic phenotypes in the fly as the locus was linked to coronary artery disease in pervious GWASs. However, to address this, we measured inflammation, which has a crucial role in CAD^70^, to further characterize cardiac phenotypes observed upon KD of genes near the CVD- and insomnia-related locus. Our study provides an important basis for future studies in more complex model systems to further characterize the roles of these genes on cardiovascular function. Moreover, our study focused on the effects of suppression of these genes on sleep and cardiac physiology. Future work will evaluate the effects of overexpressing these genes on each tissue and assess cell-autonomous and non-cell-autonomous effects that will have on sleep and cardiac function.

In summary, we report four important findings: first, *ATPSynC* is the only gene that is important cell-autonomously for both sleep and cardiac function and affects sleep non-cell-autonomously through the heart; second, *Lsn* is important for cardiac function cell-autonomously and only affects sleep non-cell-autonomously through the heart; third, *Imp* is important cell-autonomously for sleep and affects the heart only non-cell-autonomously through sleep; and fourth, we functionally identified inflammation as a mechanism connecting cardiovascular and sleep dysfunction, bidirectionally (Fig. 8). These findings advance our understanding of the association between CVD and sleep disorders and provide basis for future studies to help develop therapeutic strategies that prevent or attenuate insomnia and coincident CVD.

**Fig. 8.**
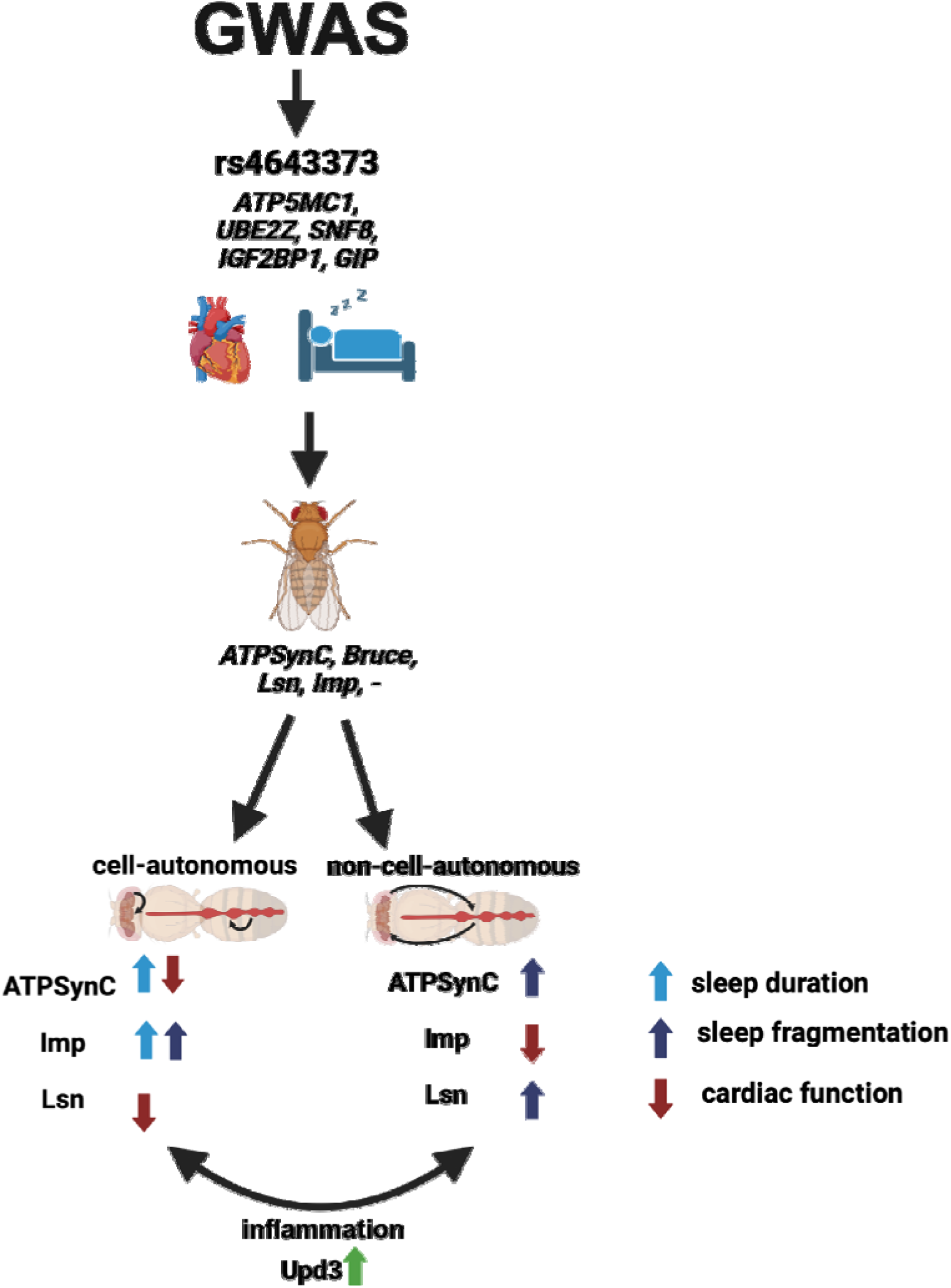
Graphical summary displaying main findings. Independent GWASs identified one genetic locus associated with CVD and insomnia. Near this locus, we identified five candidate genes, four of which have fly orthologs. To investigate the roles of genes near this locus in both neurons and the heart, we suppressed their expression in each tissue and assessed their effects on sleep and cardiac function in a cell-autonomous or non-cell-autonomous manner. Our main findings include: one gene (*ATPSynC*) involved cell-autonomously in sleep and cardiac function and affects sleep non-cell-autonomously through the heart; one gene (*Imp*) involved cell-autonomously in sleep and affects cardiac function non-cell-autonomously through sleep; and one gene (*Lsn*) involved cell-autonomously in cardiac function and affects sleep non-cell-autonomously through the heart. We also found that cell-autonomous and non-cell-autonomous effects were accompanied with inflammation through elevation of *Upd3* levels.

## Materials and Methods

### LocusZoom plots and phenotypic associations for the rs4643373 region and eQTL analyses

The LocusZoom plots for CVD and insomnia SNP, rs4643373, were generated using LocusZoom v1.3 (06/20/2014). Nearby associations to rs4643373 were obtained from the Cardiovascular Disease Knowledge Portal (broadcvdi.org); 20230927; https://cvd.hugeamp.org/variant.html?variant=rs4643373 (RRID:SCR_016536). Results were plotted using LDassoc in LDLink. The GTEx V8 database was used for multi-tissue eQTL analyses. The GTEx V8 data contains a total of 17,382 RNA-seq samples from 948 post-mortem donors. The GTEx data used for multi-tissue eQTL analyses described in this manuscript were obtained from the GTEx Portal on 03/08/2023.

### Drosophila Stocks

Drosophila stocks were cultured at 25°C on standard agar media.^71^ UAS-RNAi transgenic stocks of CVD- and insomnia-related genes were obtained from Vienna Drosophila Resource Center (VDRC) and Bloomington Drosophila Stock Center (BDSC): *ATPSynC*-RNAi (VDRC: <u>106834</u>; BDSC: **35464**, 57705), *Lsn*-RNAi (VDRC: <u>110350</u>, **21658**; BDSC: 38289), *Bruce*-RNAi (VDRC: <u>107620</u>, 48309; BDSC: **51814**), *Imp*-RNAi (VDRC: 20321, <u>20322</u>; BDSC: 38219, **55645**, 34977), control lines (*w*^1118^, VDRC: 60000, 60100, BDSC: 36303, 36304, 41552, 41553), *Act5C-Gal4* (BDSC: 4414); *24b-Gal4* (BDSC: 1767), *Elav-Gal4* (BDSC: X: 458 II: 8765). *Hand-Gal4* was obtained from Dr. Olson’s lab. Genotypes are listed in Table S4. Data from RNAi lines were not combined if more than one RNAi line was used. Line 1 is underlined and is the main line used for all experiments. Line 2 is bolded and used only in Fig. S2. *ATPSynC* Line 2 is used for neuronal-specific suppression as Line 1 was lethal when crossed with *Elav-Gal4*. *Elav-Gal4 (II)* was used in overexpression experiments to allow for observing more subtle phenotypes induced by lower expression of constructs, as overexpressing with *Elav-Gal4 (X) was lethal*.

### Ubiquitous- and Tissue-specific knock-down and genetic modulation

The GAL4-UAS system^72^ was used to drive the knockdown of CVD- and insomnia-related genes ubiquitously or tissue-specifically. Adult flies possessing UAS RNAi CVD- and insomnia-related genes were crossed to *Hand-Gal4, Elav-Gal4, Act5C-Gal4, Ubi-Gal4,* or *24b-Gal4* flies and incubated at 25°C throughout development. Adult male and female F1 progeny were separated according to sex and allowed to age, with a new food source supplied every three days prior to assays of cardiac function. Age-matched adults from *w*^1118^ (wild-type), V60100 or BL36303 (VDRC and BDSC RNAi controls) were crossed with each of the *Gal4* drivers as controls. Male and female flies were screened at 1 and 3 weeks of age for cell-autonomous assays and 3 weeks for non-cell-autonomous assays in at least 2 independent experiments. All flies were kept at 25 °C, 50% humidity in a 12 h light/12 h dark (LD) cycle.

### Sleep-wake behavioral and rhythmicity analysis

Three-to-four-day old male and female progeny of *Elav-Gal4* (cell-autonomous) and 2.5-week-old male progeny of *Hand-Gal4* (non-cell-autonomous) with control and RNAi lines of each of the four genes were collected and individual flies were loaded into glass tubes containing standard fly food (n>16). Sleep-wake behavior was recorded using the Drosophila Activity Monitor (DAM, TriKinetics inc MA, USA) system in a 12L:12D cycle at 25^°^C. Drosophila activity (or wake) is measured by infra-red beam crosses in the DAM system.^73^ To evaluate the role of CVD- and insomnia-related genes in maintaining adult rhythms, 3-4-day old male flies were loaded into glass tubes and entrained for at least 3 days to a 12h:12h LD cycle, followed by 3 days in constant darkness. Rhythmicity analysis was performed for each 3-day period in constant darkness. Data was analyzed using ClockLab and RStudio. Custom R scripts and methodology used with RStudio can be found on https://github.com/jameswalkerlab/Gill_et.al. One-way ANOVA with Dunnet’s multiple comparisons test for DAM system data was performed using GraphPad Prism. Drosophila sleep was defined by a period of at least 5 minutes of inactivity, demonstrated by zero beam breaks recorded.^74^ Average sleep per 24 hours (ZT0-ZT24) of each genotype was calculated. Five days were used for analysis of 1-week-old flies, and 3 days were used for 3-week-old fly experiments to overcome decreased viability in older flies. Sleep bouts were quantified by counting the number of periods of sleep as defined above. Sleep fragmentation was defined by either the number of 1-minute wakes or sleep bouts during a 24-hour period. Data for daytime sleep is from ZT0 to ZT12 and that for nighttime sleep is from ZT12-ZT24.

### MARGO locomotion monitoring

*Drosophila* locomotion was video monitored using the MARGO (Massively Automated Real-time GUI for Object-tracking) system for automated tracking.^46,47^ Behavior boards were prepared by the addition of standard fly food at one end. Individual 3-5-day old male and mated female flies were loaded into the behavior boards such that there was one fly per channel. Flies were video recorded for 3 days at 20°C. Collected data from days 2 and 3 were analyzed using a custom script in MatLab. Speed data was calculated by the change in location of the centroid of the tracked fly between frames. This data collected at a frame rate of 4Hz and an experimental setup such that 3.4667 pixels equal 1 millimeter. Using these values, pixels per frame were converted to millimeters per second values. Frames in which flies were immobile were excluded from speed calculations. Therefore, average speed measurements are representative of the average speed while flies are in motion. Average speed values are an average of speeds from days 2 and 3 and normalized to controls set up within the same behavior board.

### Cardiac physiological analyses of semi-intact Drosophila hearts

1-week-old male and female progeny of *Hand-Gal4* (cell-autonomous) and 2.5-week-old male progeny of *Elav-Gal4* (non-cell-autonomous) with control and RNAi lines of each of the four genes were collected, and semi-intact hearts were prepared as described (n>30).^75,76^ Direct immersion optics was used in conjunction with a digital high-speed camera (at 200 frames/sec, Hamamatsu Flash 4 camera) to record 30 second movies of beating hearts; images were captured using HC Image (Hamamatsu Corp.). Cardiac function was analyzed from the high-speed movies using semi-automatic optical heartbeat analysis (SOHA) software that quantifies heart rate, heart period, diastolic and systolic diameters, diastolic and systolic intervals, cardiac rhythmicity, fractional shortening and produced the Mechanical-mode records ^75,76^.

### Cytological Studies of adult hearts

Dissected hearts from one-week old adults were relaxed by a one-minute treatment with 5 mM EGTA in hemolymph and then fixed with 4% paraformaldehyde in PBS for 30 min as previously described ^76^. Fixed hearts were stained with anti-Pericardin antibody overnight (5ug/ml, 1:10; Developmental Biology Hybridoma Bank, University of Iowa) followed by Alexa488-phalloidin for 30 min (1:1000, U0281, Abnova), which stains F-actin containing myofibrils. Samples were then mounted with Diamond Antifade Mountant with DAPI. Confocal images were taken from a Nikon A1R HD microscope (UAB) at 10X for pericardin quantification and 20X for representative images for phalloidin staining. Quantification of pericardin area in the confocal images from three to five independent male hearts per genotype was performed by thresholding images in ImageJ, then percent area was measured.^77^

### Adult-specific KD of genes

To induce adult-specific KD, we crossed flies from each genotype with either Hand-or Elav-GeneSwitch-Gal4 and put them on RU486-containing food three-to-four days after eclosion. For crosses with *Hand-GeneSwitch-Gal4*, we used food containing 200uM of RU486, while for crosses with *Elav-GeneSwitch-Gal4*, we used food containing 500uM of RU486 to allow activation of the GeneSwitch system. We changed the food twice a week for three weeks, then, we measured cardiac and sleep physiology of three-week-old male flies.

### Viability

Adult flies (n>100, males and females) with suppression of CVD- and insomnia-related genes and controls were collected on the day of eclosion from the pupal case, designated as day zero. Approximately 30 flies were placed in each vial and transferred to a new vial every three to four days. The numbers of surviving adults were counted twice a week. The numbers of surviving adults were compared to the original number of adults collected on day zero and the percentage for each day was graphed.^78^

### Hemocyte Counts

To evaluate inflammation, fly hemolymph was collected from n>100 (per replicate, 3 biological replicates) one-week-old adult male flies with cardiac-specific suppression using *Hand-Gal4* by making an incision in the thorax of flies and centrifuging them.^79^ Hemocytes were then counted by staining the hemolymph with 1:1 Trypan blue dilution and using a hemocytometer.^79^

### Real-time quantitative PCR

Dissected male hearts (n=10-12 per biological replicate, 3 biological replicates) and heads (n=10, per biological replicate, 3 biological replicates) from 1-week-old flies was placed in the RNA lysis buffer, and flash frozen. RNA from heads was extracted using Zymo Research Quick-RNA Microprep Kit with on column DNase I digestion. RNA from hearts was extracted using the RNeasy kit (QIAGEN). Quantitative PCR was performed using SsoAdvanced Universal SYBR Green supermix (Bio Rad) in a BIO-RAD CFX Opus Real-Time PCR System. Expression was normalized with 60S ribosomal protein (RPL11). Primers for qPCR are listed below: *ATPSynC-F: GCAACAGTCGGTGTCGCT; ATPSynC-R: AGGCGAACAGCAGCAGGAA; Lsn-F: TCACCAAGGAGGACATCCTAATGG; Lsn-R: TCCGGGAATGGACTGAACTATGTA; Bruce-F: AATAGCGCTCCATCTCGACCAT; Bruce-R: ATCGACCATGCACAATGCTGT; Imp-F: AATTCGCCGACCTGGAACTCT; Imp-R: ACTCGACACCGTTCAGACCAA; Upd3-F: AGCCGGAGCGGTAACAAAA; Upd3-R: CGAGTAAGATCAGTGACCAGTTC; Rpl11-F: CGATCTGGGCATCAAGTACGA; Rpl11-R: TTGCGCTTCCTGTGGTTCAC*.

Results are presented as 2^-ΔΔCt^ values normalized to the expression of Rpl11 and control samples. All reactions were performed using biological triplicates. The means and standard error of the mean were calculated in GraphPad Prism 9 software.

### Statistical Analysis

For all quantitation except transcript levels and lifespan analyses, statistical significance was determined using one-way analysis of variance (ANOVA) followed by Dunnett’s post-hoc test to determine significance between groups for sleep and cardiac physiological parameters. Comparisons to UAS controls were calculated by 1-way ANOVA followed by Šidák’s post-hoc test to assess off-target effects of RNAi lines. Additional comparisons of sleep and cardiac parameters of flies to their respective RNAi controls in Table S5 were calculated by 1-way ANOVA followed by Šidák’s post-hoc test. For expression of transcript levels in heads, statistics were calculated by 1-way ANOVA. For expression of transcript levels in hearts, statistics were calculated by Kruskal-Wallis test and without correcting for multiple comparisons to account for variability. For overexpression experiments, statistical significance was determined using an unpaired t-test. Bar graphs show mean ± SEM. For lifespan studies, data were analyzed using the Kaplan-Meier test followed by multiple comparisons between control and experimental groups. Significance was presented using p-values on figures. All statistical analysis were performed with GraphPad Prism 9.

## Supporting information

Merge file contain mutiple figures and figures legend

## Acknowledgments

We would like to thank Dr. Philip R. Jansen for the insomnia LocusZoom plot, Dr. Harrison for supplying the UAS-Upd3 overexpression stock and Dr. Olson for supplying the Hand-Gal4 driver stock. We thank Dr. Louis Dell’Italia and Dr. Jonathan Roth for their editorial comments on the manuscript. We would also like to thank Dr. Ruan Moraes for his technical support. The Genotype-Tissue Expression (GTEx) Project was supported by the Common Fund of the Office of the Director of the National Institutes of Health, and by NCI, NHGRI, NHLBI, NIDA, NIMH, and NINDS. Stocks obtained from the Bloomington Drosophila Stock Center, and Vienna Drosophila Resource Center were used in this study. Illustrations throughout the manuscript were created with BioRender.com.

## Author Contributions

G.C.M. and F.A. designed the experiments in consultation with R.S and J.W. F.A. performed all sleep and cardiac experiments. F.A. performed analysis of cardiac physiological parameters, acquisition and analysis of cytological imaging, and qPCR experiments and analysis. T.M. performed analysis of sleep data. L.O and D.P. aided F.A. in experiments. M.M. and C.C. performed LocusZoom plots with the help of R.S. S.M. performed MARGO experiments. F.A. prepared the paper with G.C.M.’s input. All authors provided feedback on the manuscript.

## Nonstandard Abbreviations and Acronyms

CVD: Cardiovascular disease
GWAS: Genome-Wide Association Study
SNP: Single-Nucleotide Polymorphism
ESCRT: Endosomal Sorting Complexes Required for Transport
HP: Heart Period
AI: Arrhythmia Index
DI: Diastolic Interval
SI: Systolic Interval
DD: Diastolic Diameter
SD: Systolic Diameter
FS: Fractional Shortening

