## Supplementary material for "Identifying novel links between cardiovascular disease and insomnia by *Drosophila* modeling of genes from a pleiotropic GWAS locus": Merge file contain mutiple figures and figures legend

**Contents:**

Figures S1-8

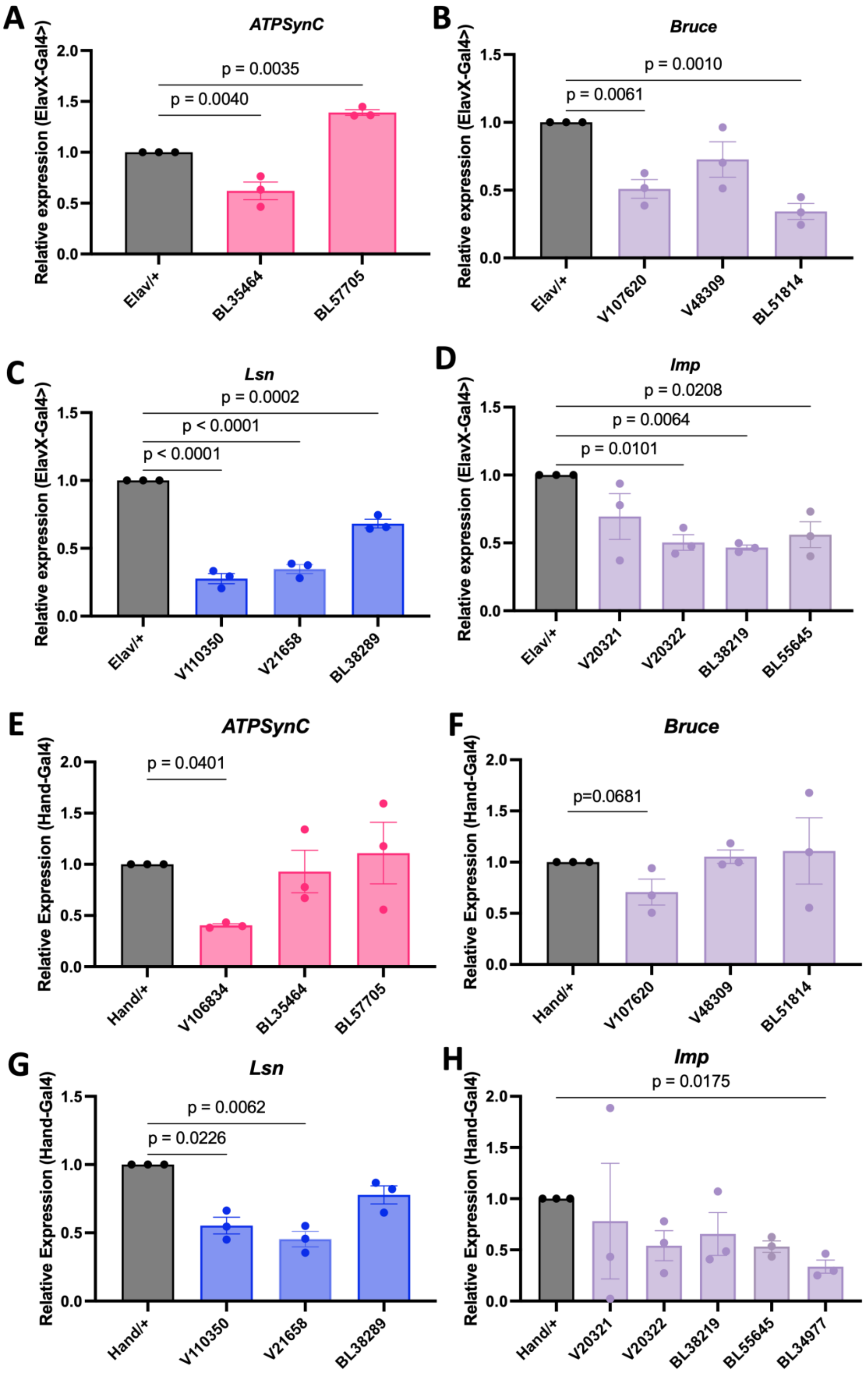

A
ATPSynC
Relative expression (ElavX-Gal4>)
p = 0.0035
p = 0.0040
Elav/+
BL35464
BL57705
B
Bruce
Relative expression (ElavX-Gal4>)
p = 0.0010
p = 0.0061
Elav/+
V107620
V48309
BL51814
C
Lsn
Relative expression (ElavX-Gal4>)
p = 0.0002
p < 0.0001
p < 0.0001
Elav/+
V110350
V21658
BL38289
D
Imp
Relative expression (ElavX-Gal4>)
p = 0.0208
p = 0.0064
p = 0.0101
Elav/+
V20321
V20322
BL38219
BL55645
E
ATPSynC
Relative Expression (Hand-Gal4)
p = 0.0401
Hand/+
V106834
BL35464
BL57705
F
Bruce
Relative Expression (Hand-Gal4)
p=0.0681
Hand/+
V107620
V48309
BL51814
G
Lsn
Relative Expression (Hand-Gal4)
p = 0.0062
p = 0.0226
Hand/+
V110350
V21658
BL38289
H
Imp
Relative Expression (Hand-Gal4)
p = 0.0175
Hand/+
V20321
V20322
BL38219
BL55645
BL34977

**Fig. S1. Transcript levels of CVD and insomnia related genes upon neuronal and CVD knockdown.** Quantification of RNA levels of ATPSynC (A), Bruce (B), Lsn (C), and Imp (D) from heads of 1-week-old male flies with following neuronal-specific suppression. Quantification of RNA levels of ATPSynC (E), Bruce (F), Lsn (G), and Imp (H) from hearts of 1-week-old male flies following cardiac-specific suppression. Each point represents 10-12 heads/hearts. Missing lines from A and D were lethal when crossed with *Elav-Gal4*. For A-D, statistics were calculated by 1-way ANOVA. For E-F, statistics were calculated by Kruskal-Wallis test and without correcting for multiple comparisons to account for variability.

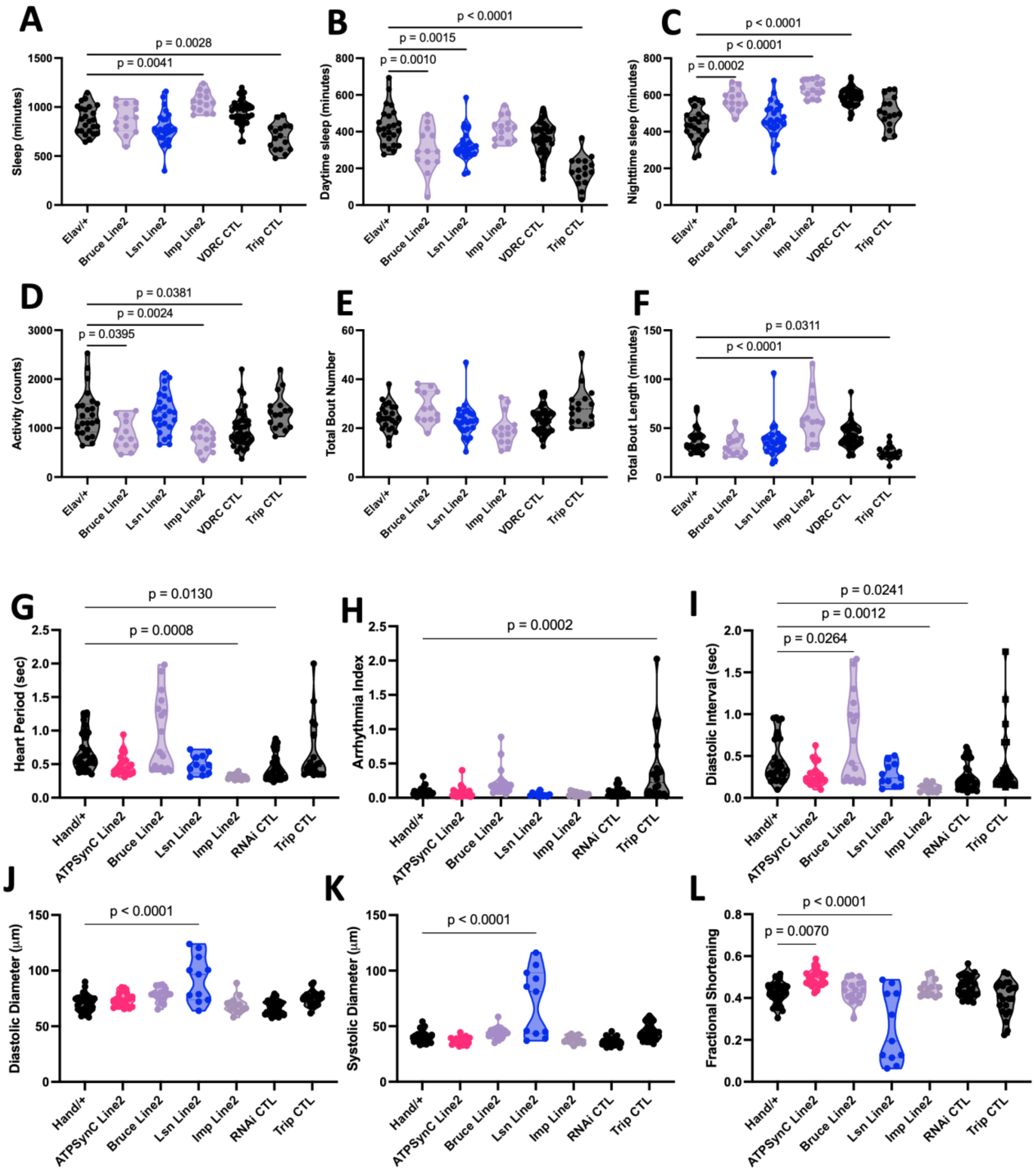


**Fig. S2. Sleep and cardiac physiological parameters of secondary RNAi line for each gene.** Violin plots for quantitative sleep parameters; sleep amount (A-C), locomotor activity (D), total bout number (E) and total bout length (F) from 1-week-old male *Drosophila* with neuronal-specific knockdown of CVD- and insomnia-related genes

(N=12-41 per group). Line 2 for each gene is: *ATPSynC* (BL35464), *Bruce* (BL51814), *Lsn* (V21658), and *Imp* (BL55645). *ATPSynC* Line 1 was lethal with *Elav-Gal4*. Violin plots for cardiac physiological parameters, heart period (G), arrythmia index (H), diastolic interval (I), diastolic diameter (J), systolic diameter (K), fractional shortening (L) from 1-week-old male flies with cardiac-specific knockdown of CVD- and insomnia-related genes (N=11-30 per group as shown in each panel, from at least 2 independent experiments). Each data point represents one fly. Statistics were calculated by 1-way ANOVA.

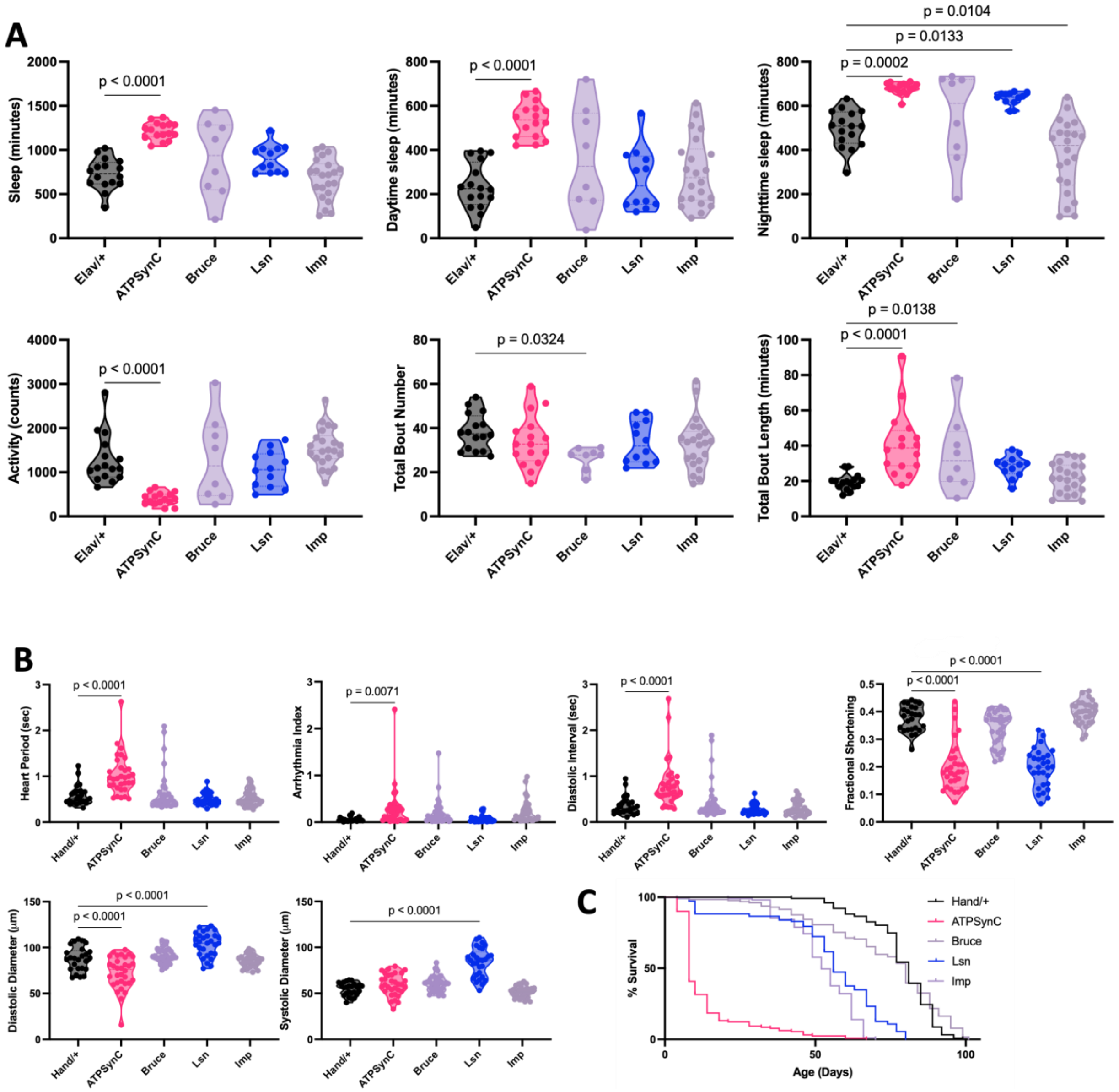


**Fig. S3. Cardiac- and neuronal-specific suppression of CVD- and insomnia-related genes leads to cardiac and sleep dysfunction in female flies.** Violin plots for quantitative sleep parameters; total sleep amount, daytime sleep, nighttime sleep, total locomotor activity, total bout number and total bout length (A) from 1-week-old female *Drosophila* with neuronal-specific knockdown of CVD- and insomnia-related genes (N=8-22 per group). Violin plots for cardiac physiological parameters, heart period, arrythmia index, diastolic interval, diastolic diameter, systolic diameter and fractional shortening (B) from 1-week-old female flies with cardiac RNAi knockdown of CVD- and insomnia-related genes with Hand-Gal4 (N=25-32 per group). Each data point represents one fly. Lifespan assay (C) for female flies with cardiac RNAi knockdown of CVD- and insomnia-related genes with *Hand-Gal4* resulted in significant decrease in lifespan ($p<0.0001$) of ATPSynC, Lsn and Imp, but a nonsignificant change of Bruce. Graph plots % survival (n>100 for each group) vs. time post-eclosion. Statistics were calculated by 1-way ANOVA for C-D and a Kaplan-Meier test was performed for G.

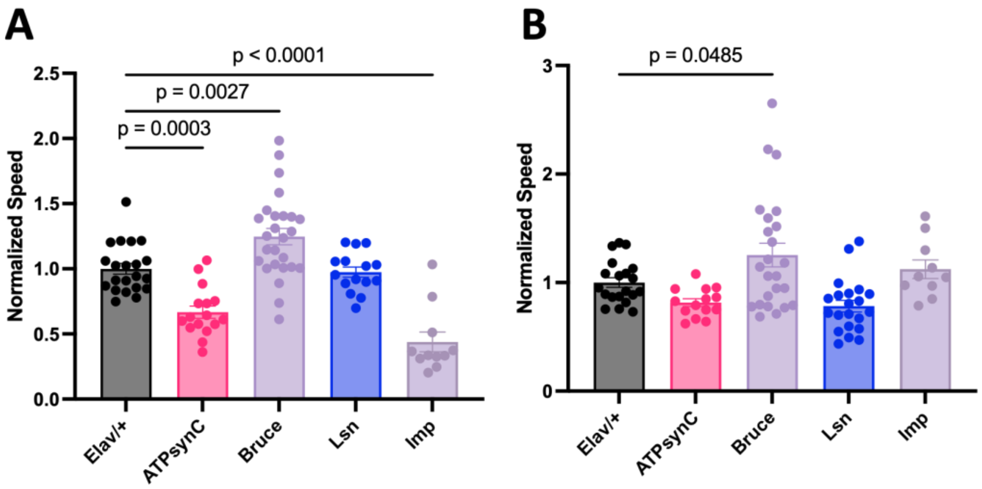


**Fig. S4. Neuronal-specific suppression of CVD- and insomnia-related genes affects locomotion speed.** Normalized locomotion speed of flies of male (A) and female (B) flies with neuronal RNAi knockdown of CVD- and insomnia-related genes with *Elav-Gal4* (n=11-26) as determined by MARGO. 1-week-old flies used. Each data point represents a fly. Statistics were calculated by one-way ANOVA for comparison to controls.

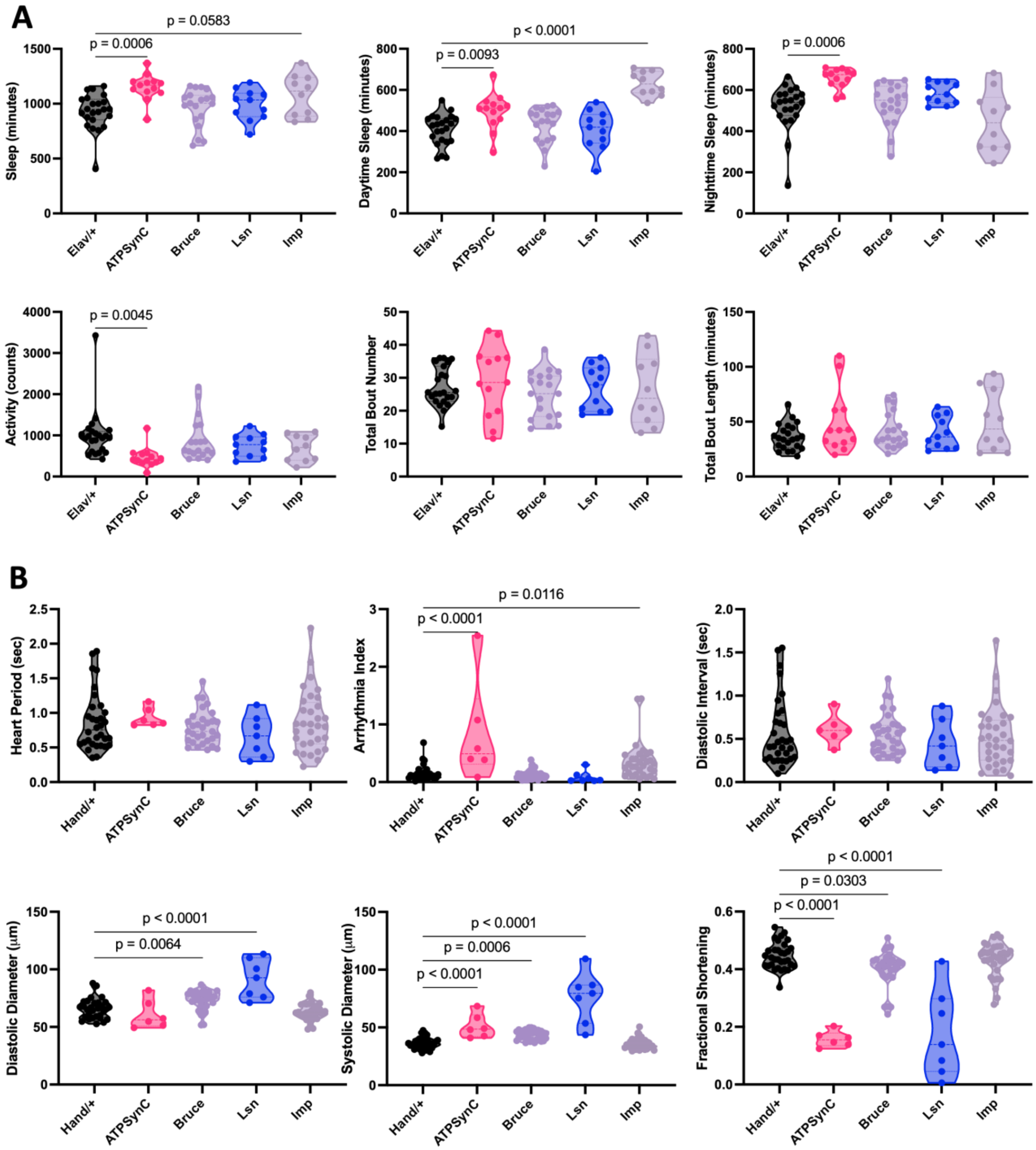


**Fig. S5. Cardiac- and neuronal-specific suppression of CVD- and insomnia-related genes leads to cardiac and sleep dysfunction in 3-week-old flies.** Violin plots for quantitative sleep parameters; total sleep amount, daytime sleep, nighttime sleep, total locomotor activity, total bout number and total bout length from 3-week-old male *Drosophila* with neuronal-specific knockdown of CVD- and insomnia-related genes (A). Violin plots for cardiac physiological parameters*,* heart period, arrythmia index, diastolic interval, diastolic diameter, systolic diameter and fractional shortening from 3-week-old male flies with cardiac RNAi knockdown of CVD- and insomnia-related genes with *Hand-Gal4* (B). Each data point represents one fly. Statistics were calculated by 1-way ANOVA.

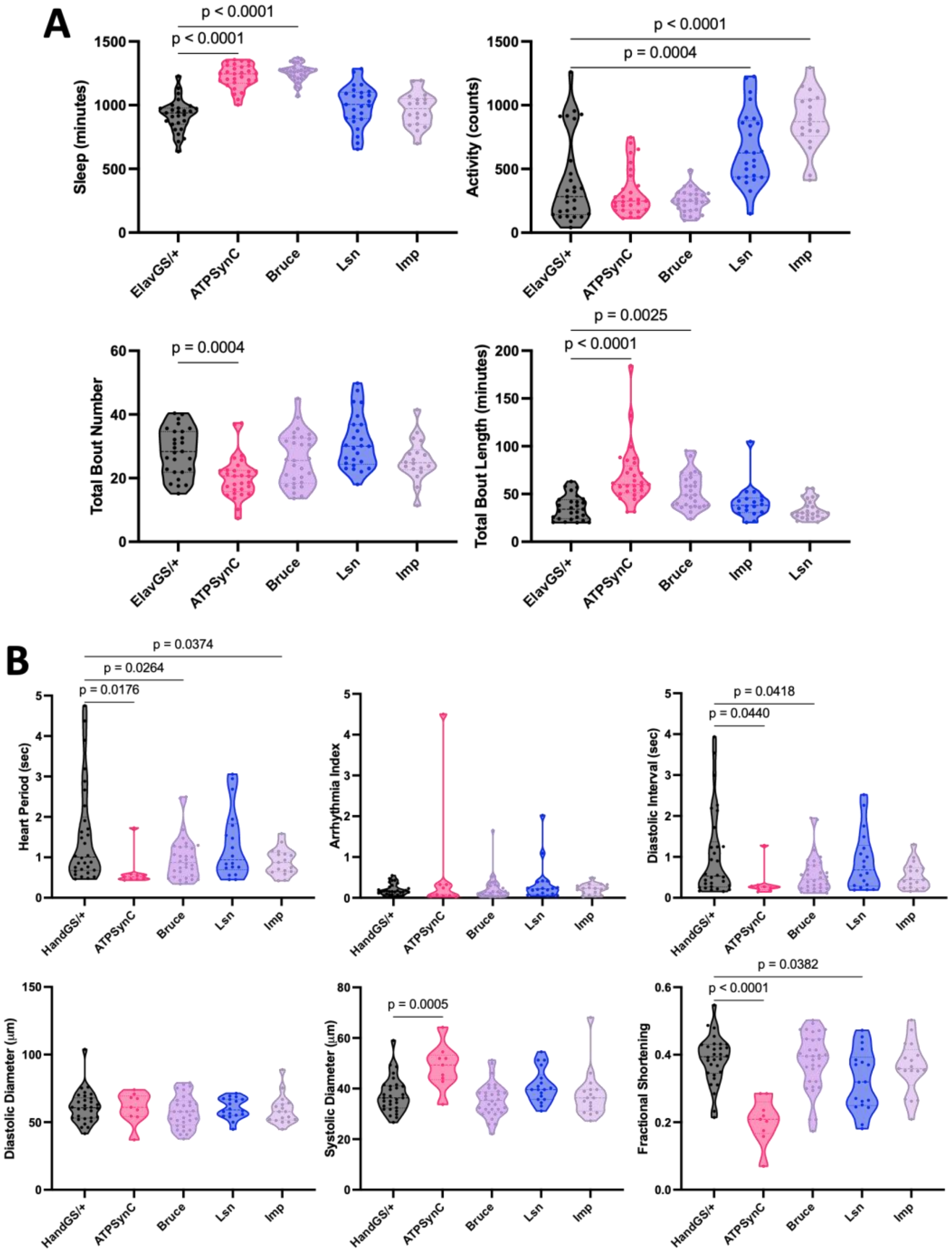


**Fig. S6. Adult-specific neuronal- and cardiac-specific suppression of CVD- and insomnia-related genes compromise sleep and cardiac function.** Violin plots for quantitative sleep parameters; sleep amount, locomotor activity, and total bout number and length (A) from 3-week-old male *Drosophila* with adult-specific neuronal-specific knockdown of CVD- and insomnia-related genes using *Elav-GeneSwitch-Gal4*. Violin plots for cardiac physiological parameters, heart period, arrythmia index, diastolic interval, diastolic diameter, systolic diameter and fractional shortening (B) from 3-week-old male flies with adult-specific cardiac RNAi knockdown of CVD- and insomnia-related genes with *Hand-GeneSwitch-Gal4.* Each data point represents one fly. Statistics were calculated by 1-way ANOVA.

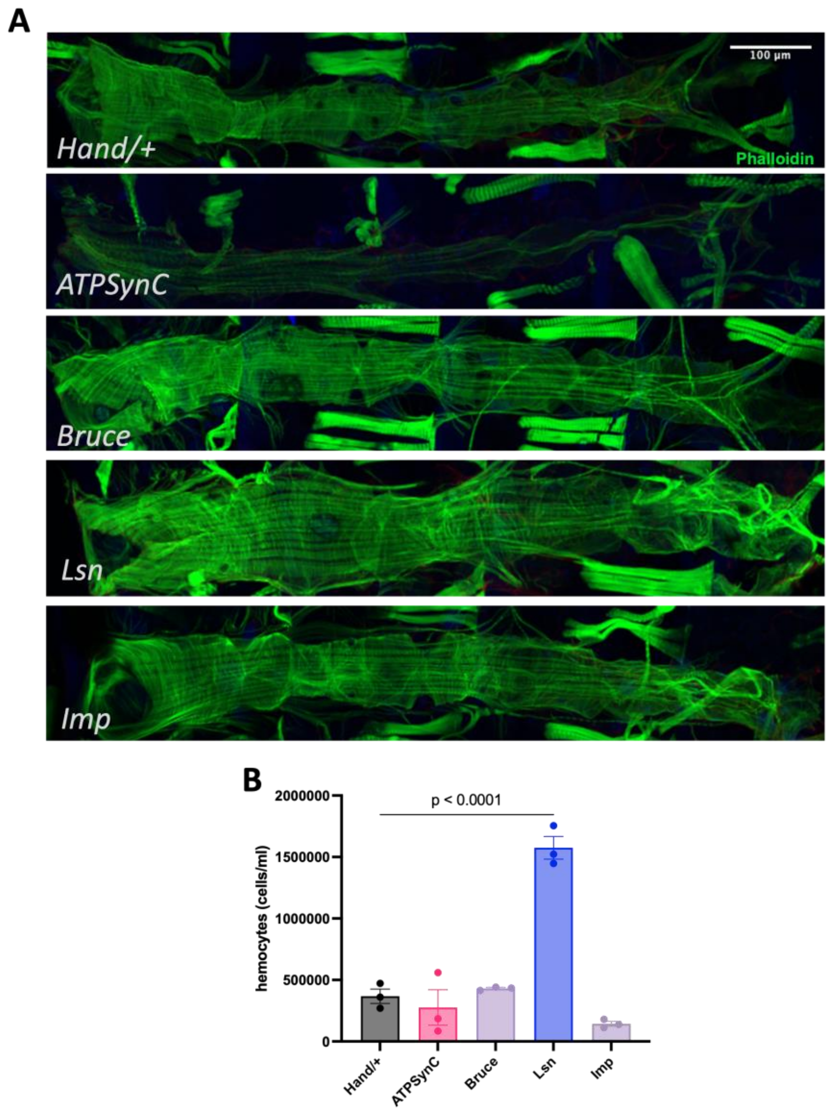


**Fig. S7. Cardiac suppression of CVD- and insomnia-related genes influences cardiac cytology and inflammation.** Representative images showing actin-containing myofibrils in whole hearts stained with Phalloidin from each group (A). Hemocyte counts (n=95-145 per data point per group) for each group (B). 1-week-old males were used. Statistics were calculated by 1-way ANOVA.

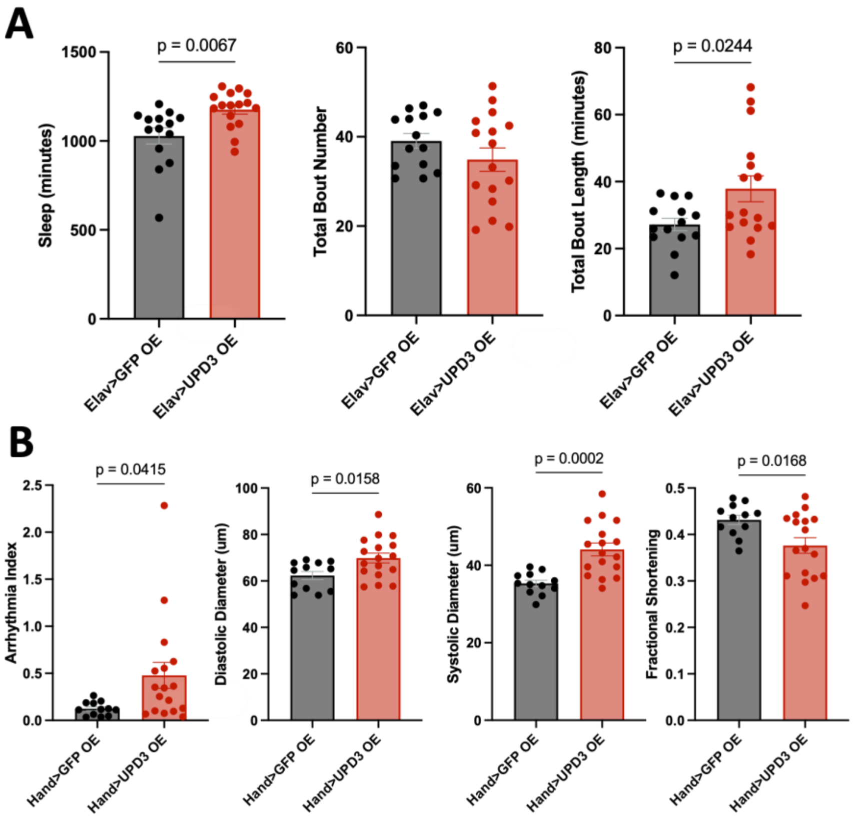


**Fig. S8. Overexpressing Upd3 in the heart leads to cardiac dysfunction, while overexpressing it in neurons does not affect sleep.** Bar graphs showing quantitative sleep parameters; total sleep amount, total bout number, and total bout length from 3-week-old male flies with neuronal-specific overexpression of Upd3 (A). Bar graphs showing cardiac physiological parameters, arrythmia index, diastolic diameter, systolic diameter and fractional shortening from 3-week-old male flies with cardiac-specific overexpression of Upd3. N= 12-16 per group. Each data point represents one fly.

Statistics were calculated by unpaired t-test.
